# Coordinated Chemokine Expression Defines Macrophage Subsets Across Tissues

**DOI:** 10.1101/2023.05.12.540435

**Authors:** Xin Li, Arlind B. Mara, Shawn Musial, Kavita Rawat, William T. King, Fred W. Kolling, Nikita Gerebtsov, Claudia V. Jakubzick

## Abstract

Tissue-resident macrophages in the lung comprising alveolar and interstitial macrophages (IMs) display a high degree of heterogeneity. In general, macrophage heterogeneity is thought to arise from various forms of activation that are heavily confounded by the recruitment of monocytes to the tissue-resident macrophage pool. To better understand the functional heterogeneity of IMs in the lung, we profiled the transcription of resident CD206^hi^ and CD206^lo^ IMs under steady-state and inflammatory conditions, excluding recruited macrophages. Rather than observing conventional *in vitro* M1 and M2 activation states, we identified seven chemokine-expressing IM subsets: IMck1 (*Ccl2, Ccl7, Ccl12,* and some *Cxcl14*), IMck2-4 (*Ccl3, Ccl4, Ccl5, Cxcl1, Cxcl2,* and *Cxcl3*), IMck5 (*Ccl8*), IMck6 (*Ccl6* and *Ccl9*), IMck7 (*Cxcl9* and *Cxcl10*), IMck8 (*Cxcl13*), and IMck9 (*Ccl24*), which were found in steady-state or induced by acute inflammation. Beyond the mouse lung, similar coordinated chemokine signatures were observed in macrophages and monocytes from other tissues and across species. Although all IMs expressed *Pf4* (CXCL4), mainly CD206^hi^ IMs were selectively depleted in *Pf4*^Cre^*R26*^EYFP-DTR^ mice. Loss of CD206^hi^ IMs resulted in significantly reduced inflammatory cell influx in allergen- and infection-driven models, as well as significantly diminished tertiary lymphoid formation and subsequent accumulation of GL7^+^ germinal center B cells. Overall, our study highlights a division of labor among interstitial macrophages, reflected by the coordinated production of chemokines to control inflammatory cell influx and organize tertiary lymphoid tissue architecture.

**One Sentence Summary:** The study highlights a division of labor among interstitial macrophages, reflected by the coordinated production of chemokines to control inflammatory cell influx and organize tertiary lymphoid tissue architecture.

## Introduction

The lungs harbor two distinct types of tissue-resident macrophages that play critical roles in maintaining homeostasis, metabolism, and tissue repair, while also functioning as sentinel phagocytic immune cells (*1, 2*). Alveolar macrophages (AMs) serve as the first line of defense in the alveoli and airways, while lung interstitial macrophages (IMs) act as gatekeepers of the vasculature and lung interstitium (*3*) (*2, 4*). Additionally, during inflammation, monocytes recruited to the lungs become part of the lung macrophage pool, known as recruited macrophages (recMacs), exhibiting their own unique transcriptional signature and function (*2, 5, 6*). Whereas we know much about the function of AMs and recruited macrophages in steady-state and in various inflammatory lung diseases, little is known about IMs.

Lung IMs are a subset of myeloid cells characterized by their expression of MerTK, CD64, F4/80, and CD11b. These IMs can be further classified into two main subsets based on the intensity of CD206 expression. The CD206^hi^ subset also expresses CD163 and Folr2, while showing low levels of CX_3_CR1. On the other hand, the CD206^lo^ subset expresses CD11c and MHCII, and exhibits higher levels of CX_3_CR1 (*7–10*). Lung IMs mainly reside in the interstitial space around the bronchovascular bundle, although a few have been found within the airspace and alveolar interstitium (*11, 12*). IMs are in close proximity to various non-hematopoietic cell types such as neurons, lymphatic vessels, and blood vessels, suggesting intensive communication and potential macrophage niche. It has been proposed that IMs provide essential trophic factors to support processes such as angiogenesis, neurogenesis, and organ development (*8, 13–20*). However, recent studies in mice lacking pulmonary IMs suggest that these cells may have additional functions beyond development and crosstalk with non-hematopoietic cells (*16*). During inflammation induced by allergens, microbes, and cancer, IMs play a critical role in regulating immunity by secreting immunoregulatory molecules, capturing and transporting antigens, and recruiting inflammatory effector cells through chemokine production (*11*). These functions have sometimes been mistakenly attributed to monocyte-derived dendritic cells (DCs) due to their shared markers with IMs (*21–23*). In the broader macrophage field, macrophage heterogeneity based on myeloid marker expression has also been linked to activation states (*24*). Classically activated macrophages, often referred to as M1 macrophages, receive instruction from Th1 cells and express markers such as inducible nitric oxide synthase (*Nos2*), IL6, CD11c, and MHCII. On the other hand, alternatively activated or M2 macrophages receive instruction from Th2 cell-produced IL-4 or IL-10 and express markers such as CD206, CD163, Arg1, Chil3, Retnla, Mgl1 and Mgl2 (*25, 26*). However, it is important to note that many of the defining markers of the M1/M2 concept, such as CD206, CD163, CD11c, and MHCII, overlap with the cell surface markers used to discriminate IM subsets. Therefore, macrophage activation should be viewed as a flexible spectrum superimposed on the backbone of IM heterogeneity, which is also evident in steady-state, with tissue- and context-specific parameters serving as dominant predictors of macrophage function (*24, 27*). Furthermore, IMs are not exclusive to the lung; but rather present throughout the body and exhibit a broadly overlapping transcriptional profile across multiple organs, suggesting that IMs have a conserved and specialized sub-tissular function, irrespective of their specific organ of residence (*7, 8*).

Here we investigated the heterogeneity and function of lung IMs in steady-state and inflammation. To gain a comprehensive understanding of IMs, we employed single-cell RNA-sequencing (scRNA-seq) to analyze IMs at a granular level by specifically enriching for IMs while excluding recently recruited macrophages and neighboring AMs. By enriching for both naïve and activated pulmonary IMs and expanding our scRNA-seq analysis to incorporate macrophages datasets derived from the airspace, pleural cavity, skin, and heart, we were able to identify and compare seven conserved chemokine signatures present in different organ-specific IM populations. Moreover, tissue-resident macrophages such as AMs and Langerhans cells also exhibited some of these conserved chemokine populations, in addition to their own distinctive chemokine signatures. Surprisingly, in IMs, we found little evidence for the production of tissue-building trophic factors and a lack of expression of the markers traditionally used to define M1 or M2 subsets, albeit there are M1 and M2 genes co-expressed in the CD206^lo^ and CD206^hi^ IMs, it is not as coordinated and defined as one observes *in vitro*. In contrast, recently recruited macrophages expressed either M1 or M2 gene profiles at a higher level. Lastly, we observed that all murine IMs expressed *Pf4* (CXCL4), which allowed us to construct a mouse model used for conditional depletion of an IM subset. Using this mouse, it was revealed that CD206^hi^ IMs orchestrate cellular influx during inflammation and organize tertiary lymphoid organs in allergen and microbial models. Overall, our findings highlight the functional diversity of IMs and their role in recruiting leukocytes and regulating immune responses in the lungs and other tissues.

## Results

### Identification and isolation of mouse pulmonary IMs

To investigate the functional heterogeneity of IMs under steady-state and inflammatory conditions, we isolated CD45^+^CD11b^+^CD206^lo->hi^ IMs from naïve mice and mice administrated with intranasal lipopolysaccharide (LPS), by enriching with CD11b beads and employing a Siglec F-Ly6G- gating strategy to eliminate AMs and neutrophils, respectively. After 24 hours of LPS treatment, recMacs become the dominant population of myeloid cells over IMs. Therefore, to capture a high number of activated IMs on the 10X scRNA-seq platform and avoid the contamination of recMacs, we depleted Ly6C^+^ monocytes by injecting the anti-Gr1 antibody before LPS treatment in one group of mice (Fig. 1A-B). An integrated UMAP analysis was performed on all three groups (Fig. 1B).

**Fig. 1.**
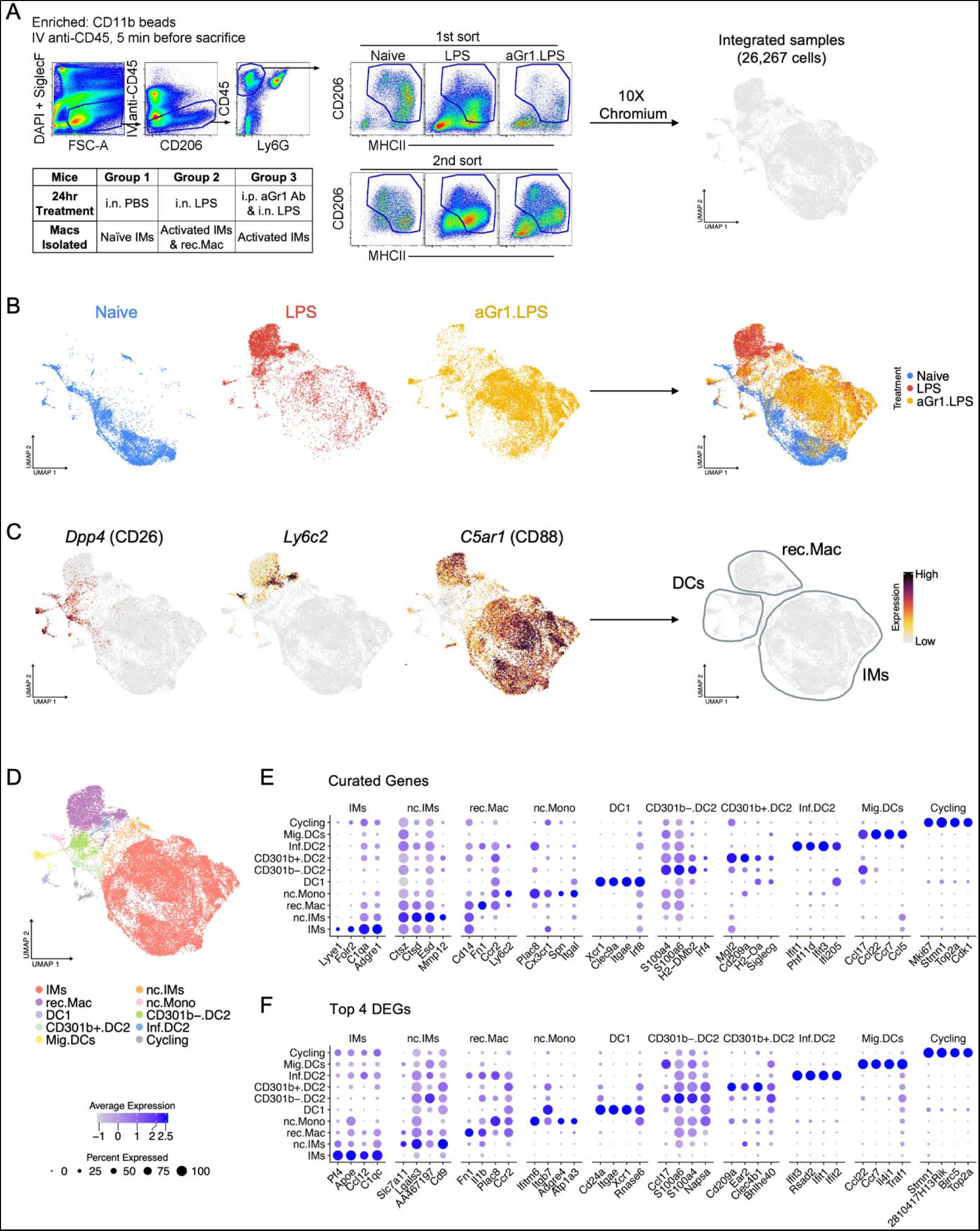
Identification and isolation of mouse pulmonary IMs. (**A**) Schematic overview of scRNA-seq experimental design, gating strategy of sorted IMs, and UMAP visualization of integrated data. (**B**) UMAP plots show the sample distribution from the individual experimental groups. (**C**) Feature plots show *Dpp4*(CD26)^+^ DCs, *Ly6c2*^+^*C5ar1*(CD88)^+^ recMacs and *Ly6C2*^-^*C5ar1*(CD88)^+^ IMs. (**D**) UMAP plot shows the distribution of 10 major cell types: IMs, non-classic IMs (nc.IMs), recruited macrophages (rec.Mac), non-classic monocytes (nc.Mono), DC1, CD301b^-^.DC2, CD301b^+^.DC2, inflammatory DC2 (Inf.DC2), migratory DCs (Mig.DCs), and cycling cells (Cycling) based on curated genes (**E**) and differentially expressed genes (DEGs) (**F**).

Sorted IMs captured a few other myeloid cell types (Fig. 1C). *Dpp4* (CD26) and *C5ar1* (CD88) were used to discriminate CD88^+^ monocyte-macrophages from CD26^+^ conventional DCs (cDCs) (Fig. 1C)(*28–30*). Within the integrated UMAP, mononuclear phagocyte clusters expressing *Ly6c2^+^C5ar1^+^* were identified as recMacs, while those expressing *Ly6c2^-^C5ar1^+^*were identified as IMs (Fig. 1C). After the identification of recMacs, IMs, and DCs, a higher-resolution analysis was performed. Ten clusters were identified using curated gene markers and top differentially expressed genes (DEGs): IMs (*C1q*, *Pf4* and *Apoe*), nonclassical IMs (expressing lower *C5ar1, Pf4* and *C1q* than all IMs), recMac (*Cd14*, *Fn1*, *Ccr2*, and *Ly6c2*), nonclassical monocytes (*Plac8*), cDC1 (*Xcr1* and *Clec9a*), cDC2 (*CD209a* and *Mgl2^hi-lo^*) that was further sub-divided into CD301b^+^ and CD301b^-^ cDC2 subsets, maturing migratory DCs (Mig.DCs)(*Ccr7*, *Ccl5*, *Ccl22*, *Il4i1*, and *Traf1*), inflammatory DCs (Inf.DCs)(*Ifit3*, and *Ifit2*), and cycling cells (*Mki67*, *Stmn1*, *Top2a*, and *Cdk1*) (Fig. 1D-F) (*11, 29*). To note, this analysis does not divide the IM subsets yet; therefore, the nonclassical IM DEGs contain similar DEGs as CD206^lo^ IMs when compared to CD206^hi^ IMs, outlined in Figure 2. Intriguingly, our analysis revealed that cDC1 and cDC2 transcriptionally converge into Mig.DCs during maturation, appearing as one cluster of cells because they lose their top signature genes while acquiring maturation genes. Mig.DCs are transcriptionally similar to mature regulatory (mreg) DCs observed in the context of cancer, highlighting the underlying mechanisms of DC maturation (*29, 31*). However, we recently published an increased granularity of Mig.DCs in the draining LN, where migratory cDC1 and cDC2 can be distinguished even in the absence of their conventional gene signatures that are no longer expressed due to maturation and migration (*29*).

**Fig. 2.**
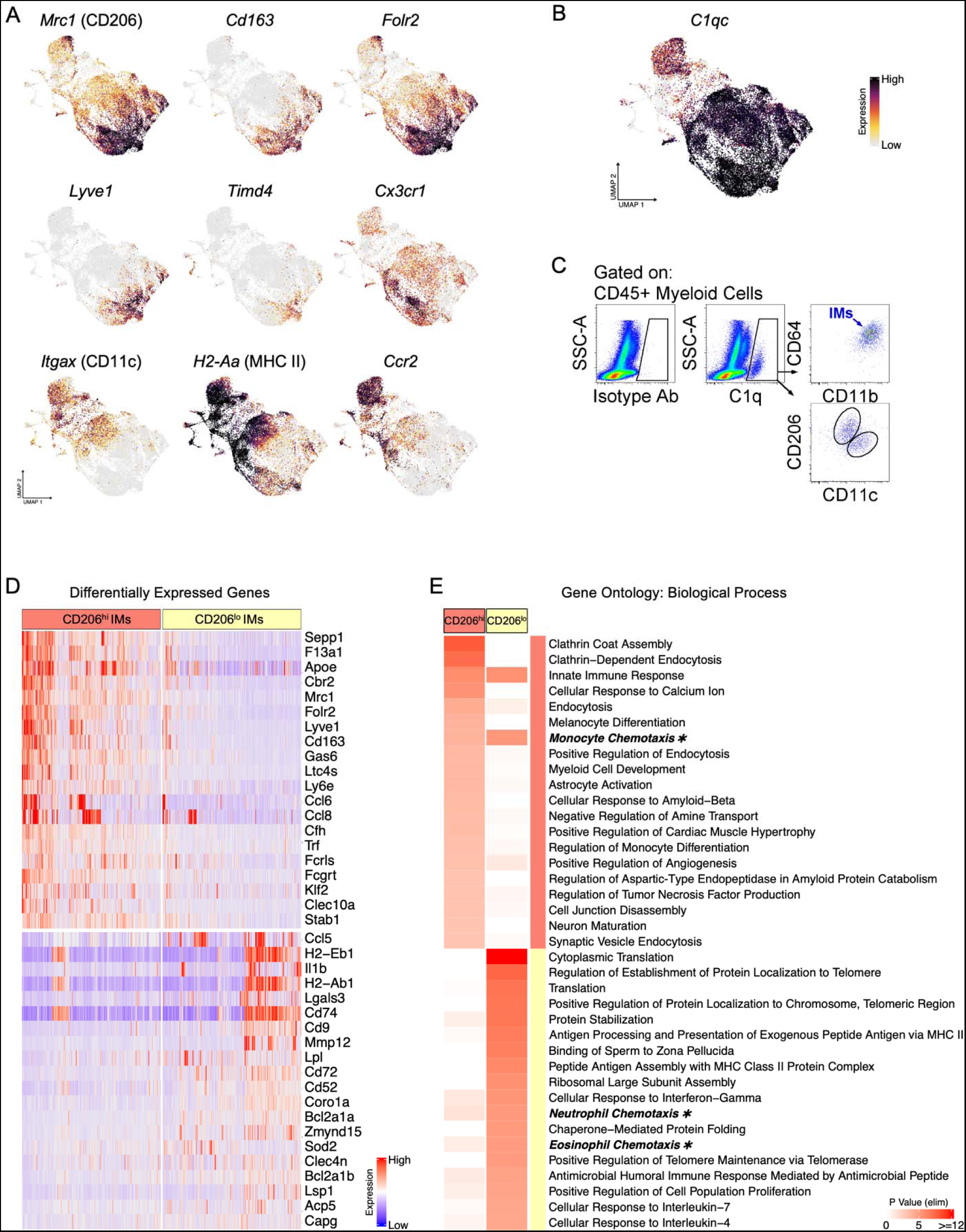
Classification of IMs into two distinct subsets: CD206^hi^CD163^hi^CX_3_CR1^lo^ IMs and CD206^lo^CD163^lo^CX_3_CR1^hi^ IMs. (**A**) Feature plots show the expression of IM genes of interest. (**B**) Feature plot shows the expression of *C1qc*. (**C**) Flow plots show the expression of C1q in IMs. (**D**) Heat map shows the expression of the top 20 DEGs in IM subsets. (**E**) Heat map shows the top 20 gene ontology terms (biological process) for each IM subset, with chemotaxis terms highlighted.

### Classification of IMs into two distinct subsets: CD206^hi^CD163^hi^CX_3_CR1^lo^ IMs and CD206^lo^CD163^lo^CX_3_CR1^hi^ IMs

IMs are classified into two main subsets based on gene and protein expression: CD206^hi^CD163^hi^CX_3_CR1^lo^ IMs and CD206C^lo^CD163^lo^CX_3_CR1^hi^ IMs (*8*). CD206^hi^ IMs express *Cd163*, *Folr2*, *Lyve1*, and *Timd4*, while CD206^lo^ IMs express *Cx3cr1*, *Itgax*, *H2- Aa,* and *Ccr2* (Fig. 2A). However, it is important to note that *Lyve1* and *Timd4* are expressed on some, and not all, CD206^hi^ macrophage subsets. Additionally, all IMs express *C1q* at both the transcriptional and protein level (Fig. 2B-C). As previously reported, DEG and Gene Ontology analyses revealed that CD206^hi^ IMs are associated with phagocytic pathways, whereas CD206^lo^ IMs are involved in intracellular protein processing and antigen presentation (Fig. 2D-E and Fig. S1) (*8, 9*). Gene Ontology analysis also suggests a role for IMs in monocyte, neutrophil, and eosinophil chemotaxis (Fig. 2E and Fig. S1F), suggesting their capacity to recruit and orchestrate leukocytes, even without further subdivision.

### Absence of unique growth factor or substantial M1/M2 gene signature expression in IM populations

Macrophages play a crucial role in maintaining homeostasis by eliminating pathogens and cellular debris. Their communication with non-hematopoietic cells through the secretion of growth factors is an essential part of this process. Given the close proximity of macrophages to various non-hematopoietic cells, such as nerves, blood vessels, and lymphatic vessels, we aimed to investigate the growth factors involved in the specific crosstalk between macrophage subsets and these cell types. Surprisingly, although some expression of growth factors was observed, our analysis did not uncover any unique growth factors expressed within unbiased clustering of the resident IM subsets. Instead, we observed broad expression of two well-known macrophage-produced fibroblast growth factors, *Tgfb1* and *Igf1* with *Igf1* being almost exclusively produced by IMs both in steady-state as well as after LPS stimulation. A few other growth factors such as *Nenf* and *Vegfb* were also produced, whereas endothelial growth factors *Vegfa* and oncostatin M (*Osm)* were mainly expressed by recMacs (Fig. S2A-B). Since growth factors did not reveal heterogeneity of IMs, we turned to M1 and M2 genes to examine potential heterogeneity (*32*). However, we did not observe broadly coordinated expression of prototypical M1 or M2 genes in IMs (Fig. S2C-D and Fig. S3). The expression of M2 markers *Psap* (prosaposin) and *Mgl2* tracked well together in baseline and after LPS stimulation of IMs, whereas on recMACs, coordinated expression was absent. Moreover, some of the most prototypical M2 genes like *Arg1* and Ym1 (*Chil3)* were almost exclusively expressed by recMacs (Fig. S3D). The few cells expressing the M1 marker *Nos2* were also almost exclusively seen in recMACs (Fig. S2C and Fig. S3A). This result is consistent with the fact that recMacs are recently recruited monocytes that enter the environment as M0 macrophages and can adopt either a pro-inflammatory (M1-like) or anti-inflammatory (M2-like) phenotype based on the needs of the environment, in addition to replenishing tissue-resident macrophage populations or becoming a monocyte-derived DCs (*33, 34*). Although IMs may seem to have relatively stable gene expression, they do not develop into *bona fide* M1-like or M2-like cells. Nonetheless, they may still exhibit pro-inflammatory and pro-resolution properties such as the expression of type 2 genes *Tslp* and *Ccl24* in skin and pulmonary CD206^hi^ IMs (*35*). Interestingly, some prototypical M2 markers like Fizz1 (*Retnla*) and Mgl1 (*Clec10a*) were coordinately expressed almost exclusively in steady-state conditions on IMs, particularly in the CD206^hi^ IM subset, potentially explaining why some investigators report high expression levels of CD206 with co-expressed markers like CD163, Folr2, and Tim4 in these IMs, considering them as indicative markers of M2 macrophages (Fig. S3C-D).

### Chemokine-expressing IM subsets are conserved across multiple organs and species

In order to further assess the heterogeneity of IMs, we performed a comprehensive and unbiased clustering analysis (Fig. S4A). The subsequent DEG analysis uncovered the enrichment of distinct chemokine genes in the different IM clusters (Fig. 3A-C and Fig. S4B). Seven chemokine-expressing subsets were identified, including IMck1 (*Ccl2, Ccl7, Ccl12,* and some *Cxcl14*), IMck2-4 (*Ccl3, Ccl4, Ccl5, Cxcl1, Cxcl2,* and *Cxcl3*), IMck5 (*Ccl8*), IMck6 (*Ccl6* and *Ccl9*), IMck7 (*Cxcl9* and *Cxcl10*), IMck8 (*Cxcl13*), and IMck9 (*Ccl24*) (Fig. 3A-I). The IMck2-4 subsets were combined since IMck4 expressed all the chemokines present in IMck2 and IMck3 (Fig. 3C). Whereas IMs expressing *Ccl24* were mainly found in samples collected from steady-state mice, while other chemokines were evidently induced by LPS exposure (Fig. 3C).

**Fig. 3.**
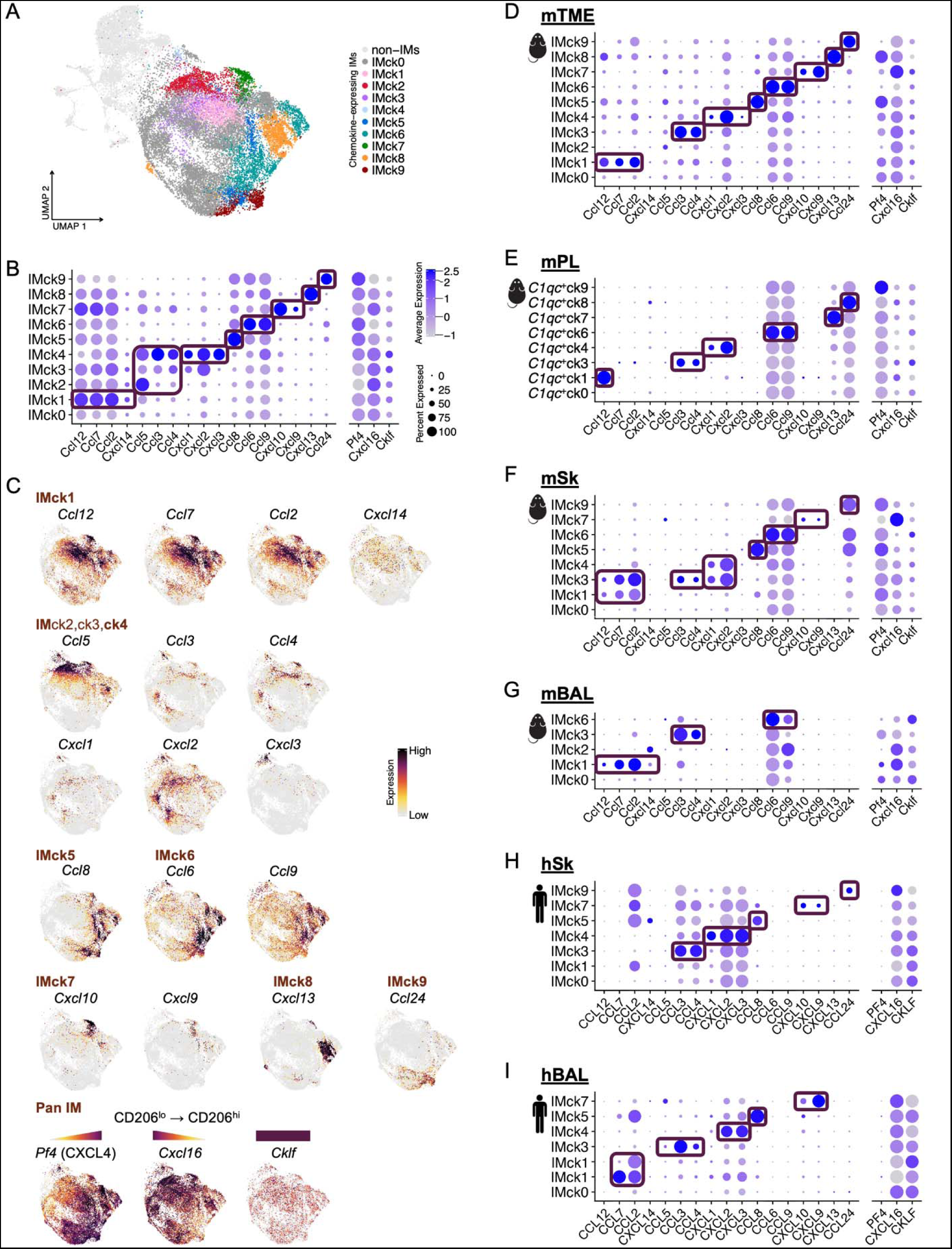
Chemokine-expressing IM subsets are conserved across multiple organs and species. (**A**) UMAP plot shows 10 IM subsets based on the expression levels of various chemokines (ck) : IMck0 to IMck9. (**B**) Dot plot illustrates the expression of chemokine genes in each IM chemokine-expressing subset. (**C**) Feature plots show the expression of chemokine genes in each IM chemokine-expressing subset. **D**-**I**. Dot plots show the expression of chemokine genes in each IM chemokine-expressing subset in different tissues and species: (**D**) Mouse tumor microenvironment (mTME); (**E**) Mouse skin (mSk); (**F**) Mouse bronchioalveolar lavage (mBAL); (**G**) Mouse peritoneal lavage (mPL); (**H**)Human skin (hSk); (**I**) Human bronchioalveolar lavage (hBAL).

Previous studies have demonstrated that, in general, IMs exhibit a conserved transcriptional profile across multiple organs and species (*7, 8*). Therefore, we investigated whether the chemokine-expressing IM subsets observed in the lungs were also present in mouse tumor microenvironments of the lung (mTME), mouse peritoneal lavage (mPL), mouse and human skin (mSk and hSk), and human bronchoalveolar lavage (hBAL) (Fig, 3D-I and Fig. S5). Interestingly, we observed the same chemokine-expressing IM subsets in all datasets within the six in-house datasets and one publicly available heart IM dataset (Fig. 3D-I and Fig. S6), indicating that these chemokine expression patterns are highly conserved. Although the chemokine genes in pulmonary IMck1 and IMck6 exhibit pan expression, their expression levels are highest within these specific clusters; where IMck1 is more selectively expressed when other IMs and C1q^+^ macrophages are analyzed across various organs and species (Fig. 3D-I). Furthermore, in lung IMs, the pan chemokine genes, *Pf4* and *Cxcl16* were inversely expressed, with *Pf4* (Cxcl4) being most highly expressed in CD206^hi^ IMs and *Cxcl16* being most highly expressed in CD206^lo^ IMs (Fig. 3C). On the other hand, the pan chemokine gene *Cklf* was expressed at similar levels in both IM subsets.

Lastly, we performed single cell trajectory analysis revealing that distinct chemokine-expressing subsets originated from either steady-state CD206^hi^ IMs or CD206^lo^ IMs, which upregulated selected chemokines upon LPS stimulation (Fig. 4). The pseudotime analysis demonstrated an enrichment of differentially expressed chemokine genes as cells progressed along the pseudotime trajectory (Fig. 4B and Fig. 4D).

**Fig. 4.**
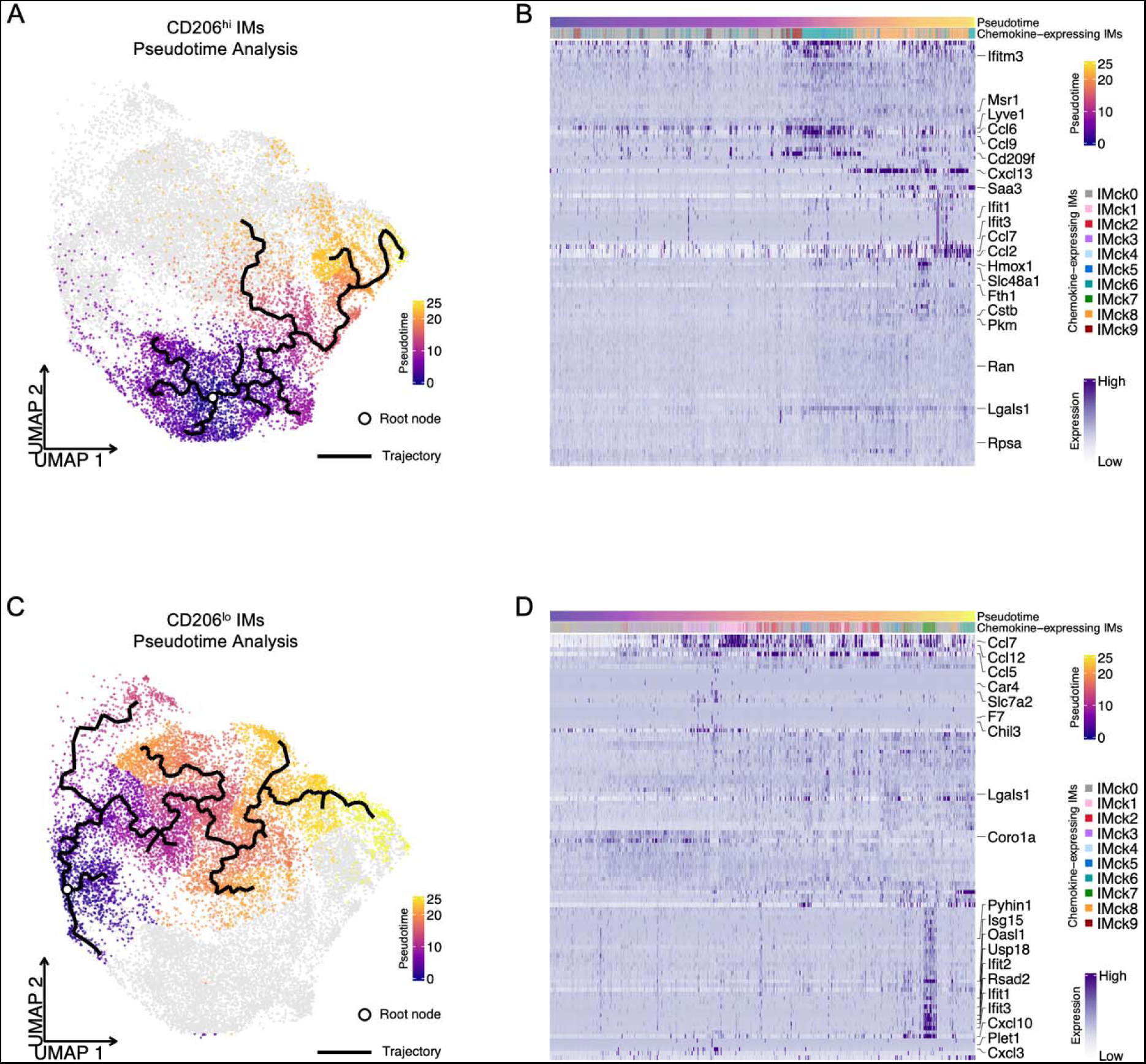
Trajectory analysis of CD206^hi^ IMs and CD206^lo^ IMs in naive and stimulated groups. (**A**) Pseudotime plot shows the trajectory of CD206^hi^ IMs from naive to stimulated states. The plot illustrates the “differentiation” of naive IMs into stimulated IMs. (**B**) Heat map shows the expression levels of the top 100 DEGs along the pseudotime trajectory across the naive-to-stimulated CD206^hi^ IMs branch. The top 20 DEGs are labeled. (**C**) Pseudotime plot shows the trajectory of CD206^lo^ IMs from naive to stimulated states. The plot illustrates the “differentiation” of naive IMs into stimulated IMs. (**D**) Heat map shows the expression levels of the top 100 DEGs along the pseudotime trajectory across the naive-to-stimulated CD206^lo^ IMs branch. The top 20 DEGs are labeled.

### SCENIC analysis identifies transcription factors associated with chemokine expression

To gain insight into the transcriptional regulation of different chemokine-expressing IM subsets, Single-Cell rEgulatory Network Inference and Clustering (SCENIC) analysis was employed, which infers the transcription factors that directly regulate the gene expression profiles based on co-expression and cis-regulatory sequences (*36, 37*). This approach yielded high specificity and accuracy, allowing us to identify putative chemokine transcription factors enriched for each chemokine-expressing IM subset, with a focus on transcription factor binding motifs located within 500 bp upstream to 100 bp downstream of the targeted gene transcription start site (TSS) (Fig. 5 and Fig. S7) or within 10k bp around the TSS (Fig. S8 and Fig. S9). Our analysis identified previously reported *Cxcl9*/*Cxcl10*-transcription factor (*38*), interferon regulatory factors, *Irf1* and *Irf9* as well as novel candidate regulators such as *Klf7* (*Ccl7*), *Xbp1* (*Ccl7*, *Ccl9*, and *Ccl2*), *Runx3* (*Cxcl16*), *Ets1* (Cxcl16), *Zeb1* (*Ccl5*), *Atf3* (*Cxcl2, Ccl4,* and *Cxcl16*), *Junb* (*Cxcl2* and *Ccl4*), *Ovol2* (*Cxcl3*), *Max* (*Cxcl10* and *Cxcl9*), *Ikzf1* (*Cxcl10*), *Stat1* (*Cxcl10, Ccl8, Ccl12, Cxcl9, Ccl5, Ccl7, Ccl4*, and *Ccl6*), *Stat2* (*Cxcl10, Ccl8, Ccl6, Ccl12, Ccl9, Ccl7,* and *Cxcl9*), and *Nfkb2* (*Cxcl10* and *Ccl4*) (Fig. 5A). We also confirmed the mRNA expression levels of these transcription factors to support the specificity of the regulon analysis (Fig. 5B). A broader range of transcription factor binding motif location (within 10k bp around the TSS) resulted in fewer enriched chemokine-regulating networks but increased specificity. Irrespective of the regulon specificity ranking, a more comprehensive analysis of all specific chemokine-regulatory networks was conducted (Fig. S7-9). The consistency between regulon specificity and transcription factor expression suggests that one could specifically target a transcription factor to regulate the differentiation of a chemokine-expressing macrophage subset.

**Fig. 5.**
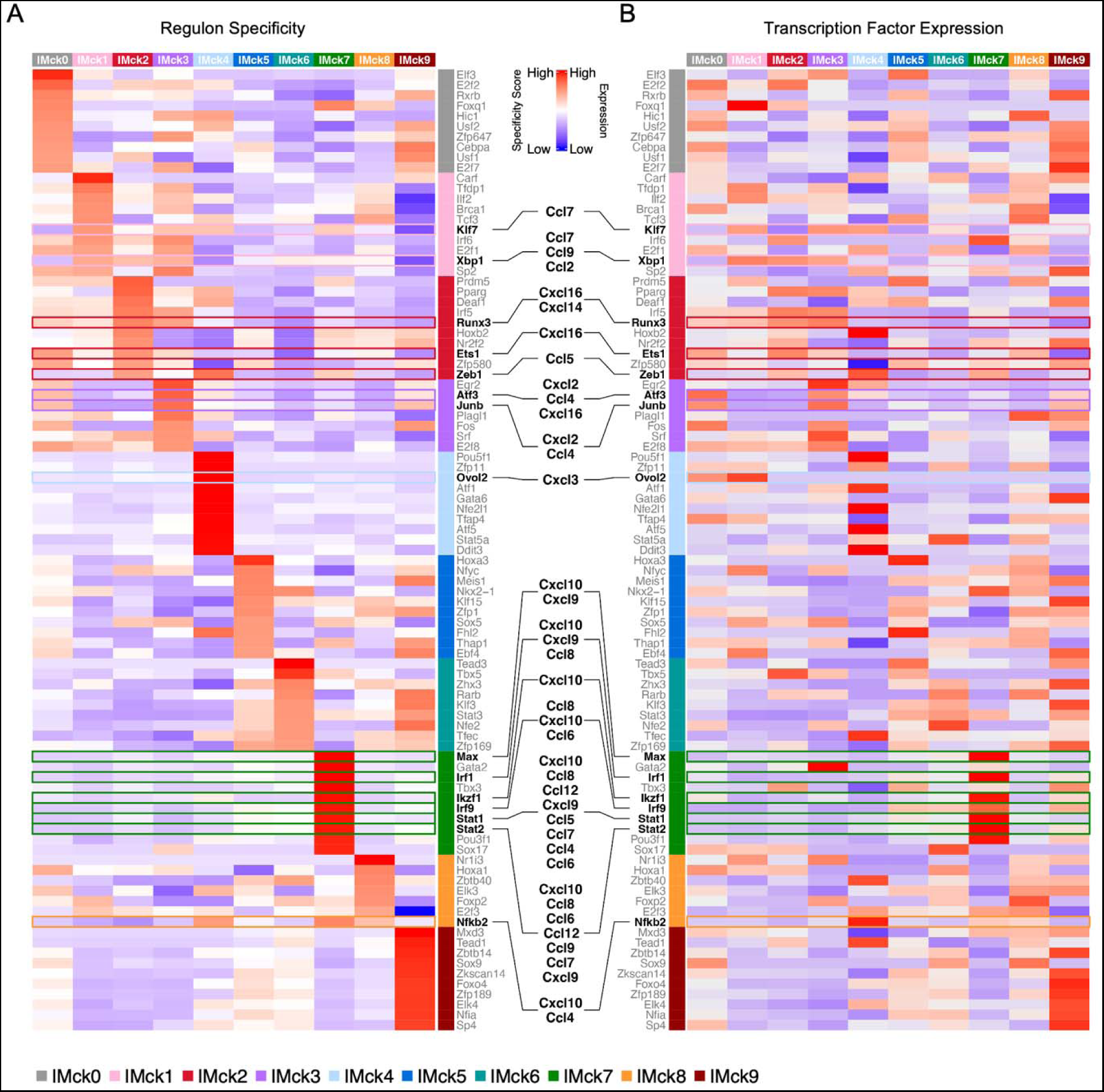
SCENIC analysis identifies regulons associated with chemokine regulation in different mouse lung IM subsets. (**A**) Heat map shows the specificity of top 10 enriched regulons in each chemokine-secreting IM subset, with the ones that putatively regulate the observed chemokine genes highlighted. (**B**) Heat map shows the expression of the transcription factors from the top 10 enriched regulons in each chemokine-secreting IM subset, with the ones that putatively regulate the observed chemokine genes highlighted.

### Tissue-specific macrophages also exhibit chemokine-expressing subsets

To investigate whether other more tissue-specific resident macrophage populations similarly express chemokines in a highly coordinated manner as observed in IMs, we analyzed large and small macrophages in the peritoneal cavity (LPMs and SPMs), alveolar macrophages (AMs) in the lungs, and Langerhans cells (LCs) in the skin of mice, as well as AMs and LCs in humans. Each of these tissue-specific macrophage populations contained some of the chemokine-expressing IM subsets and its coordinated expression pattern such as IMck1 expressing (*Ccl2* and *Ccl7*), IMck6 (*Ccl6* and *Ccl9*), and IMck7 (*CXCL9, CXCL10,* and *CXCL11*), as well as some more uniquely restricted chemokines specific to each macrophage population, such as *CCL18* in AMs or *CCL22* in LCs consistent with the idea that *CCL22* is typically known as a Mig.DC chemokine(*11, 29, 39*). Thus, not surprising to observe *CCL22* expression by LCs since LCs have DC-like migratory and antigen presenting properties that other tissue macrophages do not (Fig. S10-11). Moreover, similar analysis was performed with monocytes from different in-house scRNA-seq datasets, and similar combination of chemokine patterns were observed (Fig. S12). Overall, cross tissue and species comparison highlights the coordinated regulation of chemokine production within IMs as well as long-lived tissue-specific resident macrophages and monocytes.

### CD206^hi^ IMs contribute to iBALT formation and B cells maturation in type-2 and bacterial infection models

Since IM subsets express unique chemokine patterns, we hypothesized that they might be critical in recruiting and coordinating leukocytes during inflammation. To investigate this hypothesis, we created a mouse model that selectively depletes IMs using a single injection of diphtheria toxin (DT). Four different Cre recombinase-expressing mice were screened based on genes specifically expressed by IM subsets (Fig. 6A). All four mice were expected to eliminate at least one IM subset, if not both CD206^lo^ and CD206^hi^ IMs, when crossed with *Rosa*^LSLDTR^ (*R26*^DTR^) or *Cx3cr1*^LSLDTR^. For example, ideally crossing Lyve1^Cre^ with *Cx3cr1*^LSLDTR^ would deplete CD206^hi^Lyve1^+^ IMs while leaving Lyve1^+^ lymphatic vessels or blood vessels intact, since these cells lack the expression of CX_3_CR1. First, we wanted to ensure that Cre recombinase was expressed in the anticipated cell types by crossing *Lyve1*^Cre^, *Pdpn*^Cre^, *Nes*^Cre^, and *Pf4*^Cre^ mice with a reporter mouse, *Rosa*^LSLEYFP^ (*R26*^EYFP^). Unfortunately, *Lyve1*^Cre^ and *Pdpn*^Cre^ mice displayed a lack of Cre-specificity as EYFP was expressed in immune cells of all types rather than solely the intended cell type. On the other hand, *Nes*^Cre^ failed to express any EYFP, even though EYFP was expected to be present in CD206^lo^ IMs (Fig. 6A-B). Only the *Pf4*^cre^*R26*^EYFP^ mice exhibited Cre-specificity, where CD206^hi^ IMs selectively expressed EYFP. Based on data available on Immunological Genome Project (www.immgen.org), *Pf4* showed significantly higher expression levels in CD206^hi^ IMs when compared to CD206^lo^ IMs and other immune cell types (Fig. 6C). Although some *Pf4* expression is observed in CD206^lo^ IMs and recruited monocytes, the intensity of *Pf4*-driven Cre expression was insufficient to remove the stop cassette in the *R26*^EYFP^ mice. The ability to specifically target CD206^hi^ IMs allowed us to conduct a time-course analysis of CD206^hi^ IMs depletion using *Pf4*^Cre^*R26*^EYFP-DTR^ mice, created by crossing *Pf4*^Cre^*R26*^EYFP^ with *R26*^DTR^ (Fig. 6D). To verify the specificity of CD206^hi^ IM depletion, all macrophages were gated on, i.e., CD64^+^CD206^+^ cells. This gate identifies the three tissue-resident macrophages in the lung: auto-fluorescent CD11c^+^ AMs, CD206^hi^ IMs, and CD206^lo^ IMs. Only CD206^hi^ IMs were EYFP^+^, while CD206^lo^ IMs and AMs were not (Fig. 6D). After a single injection of diphtheria toxin (DT), CD206^hi^ IMs were depleted for up to 5 days and then gradually replenished by day 15, while CD206^lo^ IMs, AMs, and DCs remained unaffected (Fig. 6D and data not shown). With the ability to deplete CD206^hi^ IMs, their role in various diseases or inflammatory responses could be investigated.

**Fig. 6.**
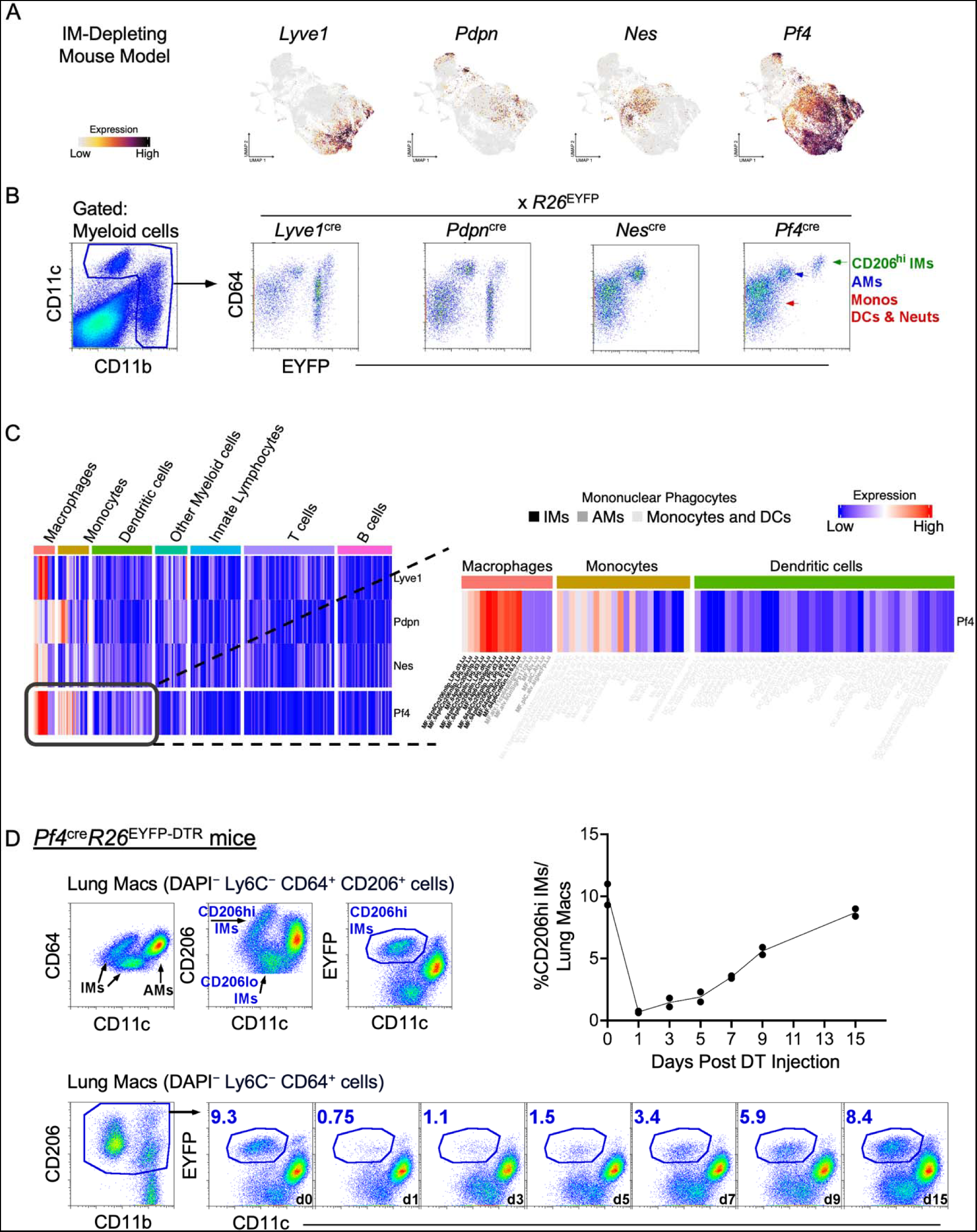
Targeting *Pf4*-expressing cells enables specific depletion of CD206^hi^ IM cells. (**A**) Feature plots show the expression of selected candidate genes for specifically targeting IMs. (**B**) Flow plots show the verification of Cre recombinase activation using *R26*^EYFP^ mice, highlighting *Pf4* as a potential marker for targeting IMs. (**C**) Heat maps show *Pf4* expression from ImmGen datasets, further supporting its selective expression in IMs. (**D**) Flow plots show gating strategy to identify CD206^hi^ IMs among all tissue-resident macrophages, followed by a time course analysis of CD206^hi^ IMs depletion after a single injection of diphtheria toxin (DT). The data presented is representative of 3 independent experiments.

Lastly, we examined the role of CD206^hi^ IMs on airway inflammation in mice sensitized and challenged with house dust mite (HDM). Previous research has shown that in models of allergen challenge, B cells aggregate to form tertiary lymphoid organs called inducible bronchus associated lymphoid tissues (iBALT) near bronchi and blood vessels (*40*). These structures contain germinal center reactions, which promote immunoglobulin class switching. In these iBALTs, B cells are thought to accumulate in a CXCL13-dependent manner with lung stromal cells as the primary source of CXCL13 (*41, 42*). However, macrophages can also be a prominent source of CXCL13 (*43, 44*). Given that CD206^hi^ IMs express *Cxcl13* (Fig. 3C), in addition to other leukocyte chemokine attractants, we studied the effect of depleting CD206^hi^ IMs during iBALT formation in the HDM model. Mice were sensitized with HDM extract. One day prior to allergen challenge, control and *Pf4*^Cre^*R26*^EYFP-DTR^ mice received a dose of DT to deplete IMs (Fig. 7A). Results showed that mice depleted of CD206^hi^ IMs exhibited significantly less peribronchial and perivascular leukocyte infiltration and iBALT formation (Fig. 7B-C and Fig. S13A). We conducted further investigation to determine whether the depletion of these cells would affect the number of mature germinal center B cells in the lungs. Indeed, the absence of CD206^hi^ IMs resulted in a significant reduction in GL7^+^CD95^+^ germinal center B cells (Fig. 7D). In addition, similar findings were observed following experimental infection with *Mycoplasma pneumoniae,* where the depletion of CD206^hi^ IMs led to reduced peribronchial and perivascular infiltrates and reduced CD95^+^GL7^+^ B cells in the lungs of challenged mice (Fig. 7E-H and Fig. S13B). Overall, these findings strongly suggest that IMs play a crucial role in leukocyte recruitment and iBALT formation in the lungs.

**Fig. 7.**
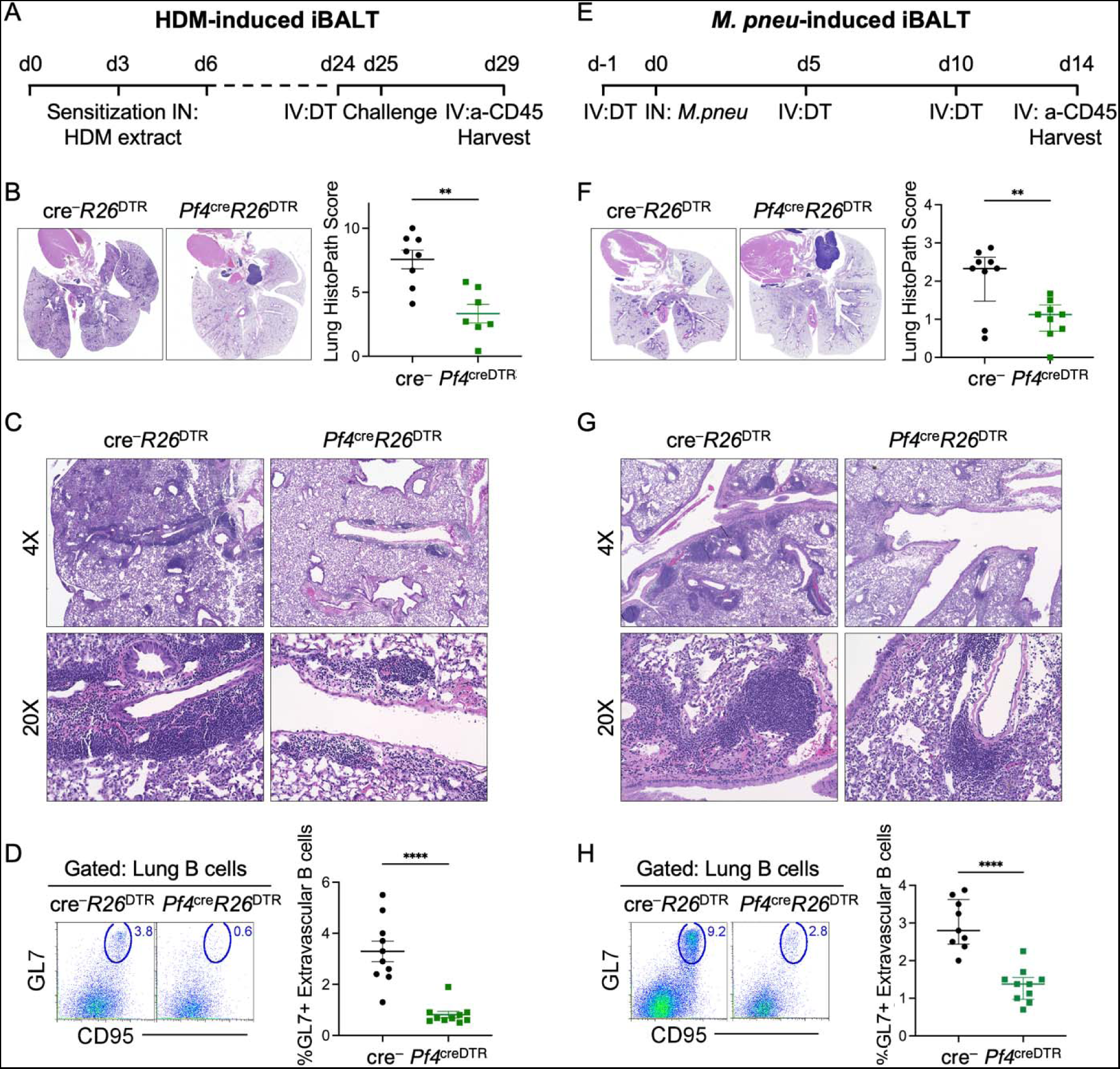
CD206^hi^ IMs contribute to iBALT formation and B cells maturation in models of type-2 and bacterial infections. (**A**) Experimental design of house dust mite (HDM) model. **B**-**C**. Representative IHC images and statistical analysis of lung histopathological score for IM depleted mice in the HDM model compared to control. Data represented are from 2 independent experiments with n=4/5 per group. (**D**) Gating strategy and statistical analysis for mature GL7^+^CD95^+^ B cells in the HDM model compared to control. Data represented are from 2 independent experiments with n=4/5 per group. (**E**) Experimental design of *Mycoplasma pneumoniae* (*M. pneu*) model. **F**-**G**. Representative IHC images and statistical analysis of lung histopathological score for IM depleted mice in the *M. pneu* model compared to control. Data represented are from 2 independent experiments with n=4/5 per group. (**H**) Gating strategy and statistical analysis for mature GL7^+^CD95^+^ B cells in the *M. pneu* model compared to control. Data represented are from 2 independent experiments with n=4/5 per group.

## Discussion

Within the pulmonary mononuclear phagocyte system, there are various cell types, including AMs, DCs, extravascular monocytes, and IMs. Our study aimed to investigate the role of pulmonary IMs during steady-state and acute inflammatory conditions. However, during inflammation, monocytes migrate into the lung giving rise to recMacs, which become the overabundant macrophage compared to tissue-resident IMs. To ensure that we focused on activated tissue-resident IMs, anti-Gr1 antibody was used to deplete circulating Ly6C^hi^ monocytes. This approach enabled us to stimulate IMs, *in vivo*, in their natural environment, rather than extracting and stimulating them *ex vivo*, where their survival is compromised. In our scRNA-seq dataset, the enriched IM population, which excluded alveolar macrophages using Siglec F in a dump channel, identified ten distinct mononuclear phagocyte populations. These included CD206^hi^ IMs, CD206^lo^ IMs, non-classical IMs, recMacs (i.e., derived from classical monocytes), non-classical monocytes, cDC1, cDC2 (CD301b^+^ and CD301b^-^ cDC2), inflammatory DC2, migratory DCs, and cycling cells. Notably, since IMs were enriched, IMs were the most abundant population captured on the scRNA-seq 10X platform.

We conducted a higher-resolution analysis of the IM population and discovered that these cells were subdivided based on the chemokines they express rather than growth factors or M1/M2 dichotomy. This finding led us to propose that IMs play a crucial role in orchestrating leukocyte recruitment during inflammation. To explore this hypothesis, we developed a mouse model that selectively depletes CD206^hi^ IMs, which revealed that IMs are indispensable for organizing and recruiting perivascular and peribronchial leukocytes in two different inflammatory models. However, it’s important to note that our findings do not negate the possibility that IMs also perform additional bone-fide macrophage functions, including crosstalk with structural cells, clearance of pathogens and cellular debris, and activation of effector T cells and other immune cells such as innate lymphoid cells and eosinophils (*35*).

While our IM-depleting mouse model provides a method to deplete a subset of IMs, our dataset, in conjunction with a recently published dataset (*45*), could potentially help other investigators generate a more specific CD206^hi^ or CD206^lo^ IM-depleting mouse models. For example, CD206^lo^ IMs express Cadm1 exclusively. Cadm1 is not expressed on microglia, CD206^hi^ IMs or recMacs. Moreover, cDC1 does not express CX_3_CR1, so if Cadm1^Cre^ were crossed with CX_3_CR1^LSLDTR^ mice, theoretically, DT treatment would selectively deplete CD206^lo^ IMs. However, this Cre mouse needs to be developed as it is not commercially available. Similarly, *Cx3cr1*^LSLDTR^ mice crossed with *Tmem119*^Cre^ could potentially target CD206^lo^ IMs and some recMacs, but BM chimera of these mice into WT mice would be required to avoid microglia depletion, which express both Tmem119 and CX_3_CR1 at high levels. However, a recent study has developed a Tmem119^cre^ mouse that selectively targets all IMs but not microglial cells, which is interesting as there is little to no expression of Tmem119 in CD206^hi^ IMs (https://cells.ucsc.edu/?ds=lung-interstitial-macrophage&gene=Tmem119) (*10*). On the other hand, CD163^Cre^, Folr2^Cre^ or CCL24^cre^ may also be appropriate for targeting all or some CD206^hi^ IMs, with the latter mouse recently developed (*35*). Nonetheless, regardless of the mouse model designed, the specificity of Cre-expression must be verified. Since, as observed, three out of the four Cre-expressing mice we hypothesized to be relatively IM specific did not demonstrate the anticipated outcome of cellular specificity. Even in the *Pf4*^Cre^ mice, although CD206^lo^ IMs and recMacs expressed *Pf4*, the Cre expression level in these cells was insufficient to remove the stop cassette in *R26*^LSLEYFP^ or R26^LSLDTR^ mice. As we continue to analyze the data, more IM or recMac- specific mouse models will be developed to enhance depletion specificity.

In the future, we aim to investigate how chemokine-expressing IM subsets are regulated and what inflammatory settings dictates their outcome. Our experiments using *Pf4*^Cre^*R26*^DTR^ mice to eliminate Cxcl13-expressing IMs and other chemokine-expressing IMs demonstrated a significant reduction in iBALT formation, highlighting the contribution of IMs in tissue lymphoid recruitment and development. However, the most intriguing discovery was that specific chemokine expression patterns in macrophages are conserved across organs and species. While some combinations of chemokines were shared with tissue-specific macrophage populations, these macrophages also expressed their unique chemokine patterns.

An interesting parallel can be drawn to T cell biology, where naive differentiated CD4 helper T cell (Th) subsets express a unique combination of cytokines and chemokine receptors, which is determined by a master transcription factor (Fig. S14). Based on our findings, we propose a model in which steady-state macrophages, like naive T cells, differentiate into distinct subsets that express specific chemokines in response to environmental stimuli. This differentiation is guided by particular transcription factors, leading to the unique set of chemokines observed in each subset (Fig. 8). Just as the identification of T-bet, GATA-3, RORγt, Bcl-6, and Foxp3 provided insight into the induction of Th1, Th2, Th17, T follicular helper cells, and Tregs from naïve T cells, identifying the master regulators in macrophages that dictate the outcome of chemokine-expressing macrophage subsets will be equally fascinating.

**Fig. 8.**
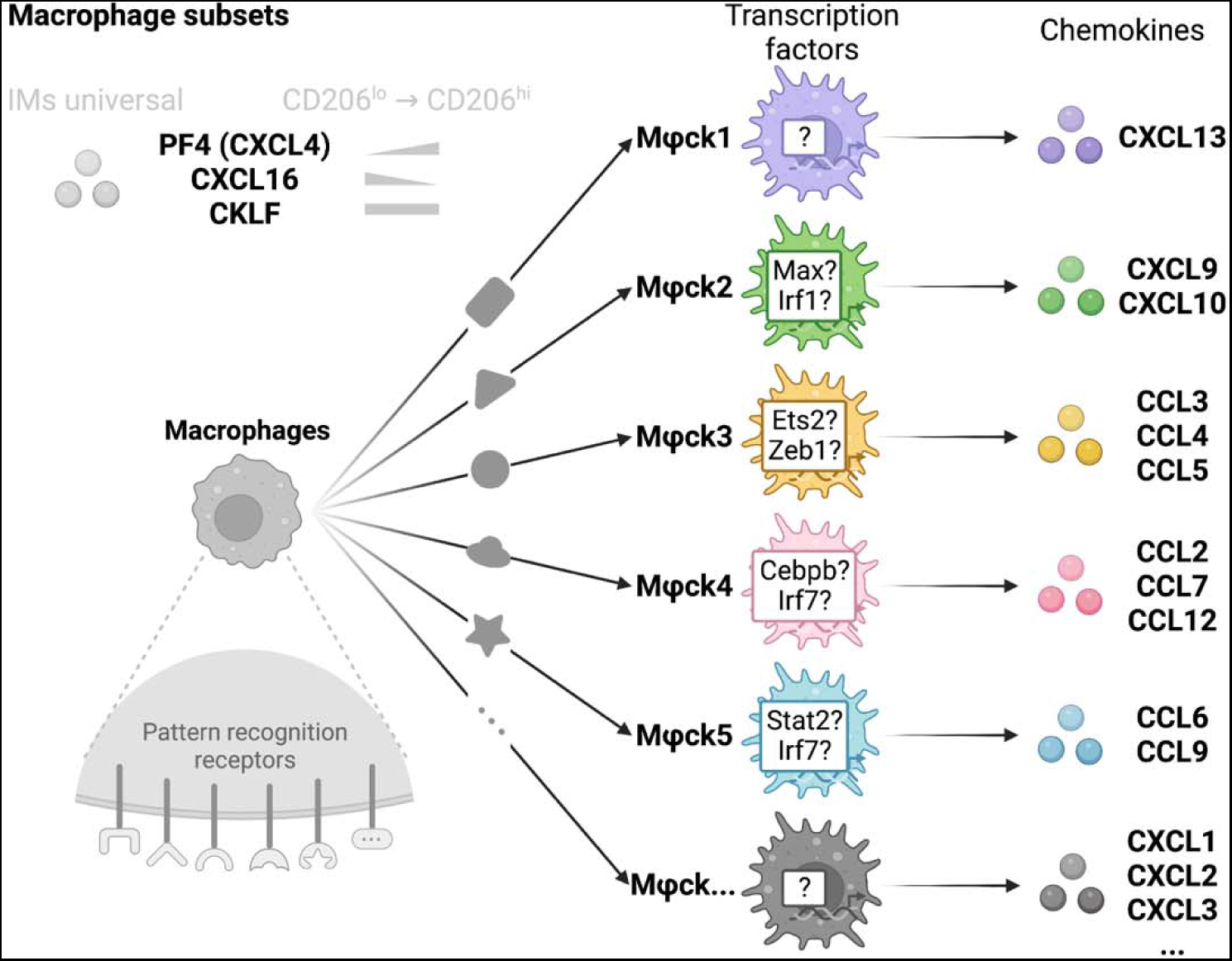
Hypothesized regulation of chemokine expression by macrophages. The diagram summarizes the proposed universal macrophage subtype biology, highlighting the specific differentiation routes and chemokine expression patterns of different macrophage subtypes.

## Material and Methods

### Mice

C57BL/6 Ly5.1 (CD45.1) WT mice were purchased from Charles River/NCI. B6;129P2- Lyve1tm1.1(EGFP/cre)Cys/J (*Lyve1*^Cre^), B6.Cg-Tg(Nes-cre)1Kln/J (*Nes*^Cre^), Tg(Pdpn- cre/GFP)1Gco/J (*Pdpn*^Cre^), C57BL/6-Tg(Pf4-icre)Q3Rsko/J (*Pf4*^Cre^), B6.129X1- Gt(ROSA)26Sortm1(EYFP)Cos/J (ROSA^fl/flEYFP^), and C57BL/6- Gt(ROSA)26Sortm1(HBEGF)Awai/J (ROSA^fl/flDTR^) mice were purchased from Jackson Research Laboratories. All mice were bred in-house. Mice were genotyped or phenotyped prior to studies and used at 6-12 weeks of age; housed in a specific pathogen-free environment at Dartmouth Hitchcock Medical College, an American Association for the Accreditation of Laboratory Animal Care accredited institution; and used under protocols approved by the Institutional Animal Care and Utilization Committee.

### Bone Marrow Chimeras

Eight-week-old CD45.1 WT mice were lethally irradiated with two rounds of 460 rads and reconstituted intravenously with freshly isolated bone marrow (BM) cells. CD45.2 Cre^+^DTR^+^ or Cre^-^DTR^+^ BM cells were prepared by flushing donor femurs and tibias with PBS. Cells were counted and transferred into recipient mice after radiation exposure. Five weeks post-reconstitution, mice were assessed for chimerism prior to use.

### Preparation of Single-Cell Suspensions from Organs

Blood was drawn by cardiac puncture using a 26-gauge syringe coated with 100 mM EDTA to prevent coagulation, followed by lysis with BD Pharm lyse (BD Biosciences, San Jose, CA) red blood cell lysis buffer. Bronchoalveolar lavage was obtained by flushing the airways four times with 1 ml of cold PBS only. Mice were perfused via the heart with 20 ml of PBS. Lung tissues were finely minced with scissors before digestion. Lungs were digested with 500 μg/ml Liberase (Roche/Sigma, Branford, CT/St. Louis, MO) for 30 minutes at 37°C. A total of 100 ml of 100 mM EDTA was added to stop 1 ml of enzymatic digestion. Single cell suspensions were obtained by repeated glass pipetting before cells were filtered through a 100-μm nylon filter. In some instances, cells of interest were enriched by MACS bead purification (Miltenyi, San Diego, CA) using anti-CD11b microbeads before analysis. To differentiate intravascular and extravascular leukocytes, mice were injected intravenously with APC-Cy7-conjugated anti-CD45 antibody 5 minutes before organ harvest.

### Flow Cytometry

Single-cell suspensions were resuspended in FACS buffer containing HBSS with 0.3 μM EDTA and 0.2% FBS, and stained for 30 minutes with antibody mastermix. Table E1 in the online supplement outlines the complete list of antibodies used. The viability dye DAPI (#D9542; Sigma) was added immediately before each sample acquisition on a BD Symphony A3 analyzer (BD Biosciences). Data were analyzed using FlowJo (Tree Star, Ashland, OR). Ag-specific Abs and isotype controls were obtained from BioLegend, eBioscience, and BD Biosciences.

### Mouse inflammatory models

#### Mycoplasma pneumoniae

*Low passage M. pneumoniae s*train PI1428 was obtained from ATCC and cultured using ATCC Medium 243 supplemented with 20% Heat-inactivated Horse Serum and 10% ATCC Yeast Extract #3. *M. pneumoniae* was cultured to high titers and frozen back in 500uL aliquots containing 5.93×10^9^ CCU viable bacteria as determined by the Most Probable Number method. On the day of challenge, frozen aliquots were resuspended in fresh medium and incubated at 37C with gentle shaking for 5 hours, pelleted, then resuspended at the target concentration. Mice were anesthetized with Avertin and intranasally instilled with 50uL containing of 5×10^8^ CCU per 50uL of medium. The inoculum was verified to contain viable titers to the 10^8^ CCU through assessing growth following 10-fold serial dilutions.

#### House Dust Mite Extract

Mice were anesthetized with aerosolized isoflurane and instilled with 25ug of *Dermatophagoide pteronyssinus* extract (Greer Laboratories) in 50uL of PBS. Experimental timelines are illustrated within the data figures.

### Microscopy

Harvested lungs were gently perfused with 5mL PBS, inflated with 10% Neutral buffered formalin in 10% neutral buffered formalin. Lung tissue was dehydrated in ethanol gradients, cleared in xylene then paraffin embedded.. Paraffin–embedded sections (5 μm in thickness) were stained with hematoxylin and eosin (H&E) for standard histology. Images were collected with a Keyence BZ-X800 Series (Keyence). The images were obtained with a 20x objective for a final magnification of x200. For Histopathological Evaluations, the sections were blindly scored based on the severity of the peribronchiolar and perivascular infiltrates which in itself is based on the size, cellularity, and organization of the lesions. A score of 0 indicates no infiltrates/lesions, 1 indicates mild infiltrates/lesions (abnormal infiltrates, often with interrupted collar), 2 indicates moderate infiltrates/lesions with complete or crescent collar with <5 cells thickness, 3 indicates marked infiltrates with crescent collar between 5-10 cells thick, and 4 indicates severe infiltrates with complete collars and 5-10 cells thick. Frequency of lesions is also taken into consideration when assigning scores, and half-step intervals are used for lesions severities meeting criteria between two scores. Each visible lobe is scored separately and the average is taken to obtain the HistoPath Score reported per specimen.

### scRNA-seq analysis of mouse lung

#### Mouse treatment

Three groups of C57BL/6 mice (n=2) were treated for cell collection. LPS was intranasally administrated to mice with 10 ug in 50 ul sterile PBS 24 hours before harvest. a-Gr1 Ab was intraperitoneally administrated to mice with 300 ug in 300 ul also 18 hours before harvest. Mice in group 1 are naïve mice without any treatment, where the collected cells are expected to contain only steady-state IMs; mice in group 2 are treated with LPS i.n. to introduce acute inflammatory lung injury, where the collected cells are expected to contain both stimulated IMs and recruited monocytes; mice in group 3 are treated with LPS i.n. and a-Gr1 Ab to introduce injury and block the recruitment of blood monocytes, where the collected cells are expected to contain only stimulated IMs. To differentiate intravascular and extravascular leukocytes, mice were injected intravenously with APC-Cy7-conjugated anti-CD45 antibody 5 minutes before organ harvest. The mice were euthanized using CO2 inhalation.

#### Sample processing

IMs were isolated from lung as previously described. Briefly, the lungs were perfused and finely minced with scissors and digested for 30 minutes at 37°C. Single cell suspensions were obtained by repeated glass pipetting and filtration. Cells were centrifugated at 300g for 5 min at 4°C and stained with fluorescent labeled antibodies. Stained single cell suspensions were enriched by MACS bead purification (Miltenyi, San Diego, CA) using anti-CD11b microbeads and then FACS sorted using the FACS Aria Fusion (BD Biosciences; strategy shown in Fig. A1). Sorted cells were centrifugated at 300g for 5 min at 4°C. Supernatants were aspirated, and pellets were resuspended with HBSS plus 0.5% BSA at an approximate concentration of 2.5 × 105 cells/ml. Cell quality and viability were assessed with a Cellometer K2 (Nexcelom Bioscience). All samples had viability >90%. Single cells were then processed using the Chromium Next GEM Single Cell 3′ Platform (10X Genomics). Approximately 6,000 cells were loaded on each channel with an average recovery rate of 5,000 cells. Libraries were sequenced on NextSeq 500/550 (Illumina) with an average sequencing depth of 30,000 reads/cell.

#### Data preparation

Raw sequencing reads were demultiplexed, mapped to the GRCm38 mouse reference genome, and gene expression matrices were generated using CellRanger v6.1 (10X Genomics). The following analyses were conducted in R 4.2 (*46*) and Python 3.6. Seurat package v4.3 was used for downstream data analyses (*47*), and figures were produced using the package ggplot2. Following a standard workflow, the gene expression matrix was filtered to discard cells with less than 200 genes, as well as genes that were expressed in less than three cells. Samples then underwent quality control to remove cells with either too many or too few expressed genes (average around 1,400 and 4,600) and cells with too many mtRNA (average around 4%), resulting in a total of 26,267 cells. Then, “SCTransform” was applied with the “glmGamPoi” method to normalize gene expression data (*48, 49*). After individual preparation, all the samples were merged into a combined Seurat object. Then scaled values of variable genes were then subject to principal component analysis for linear dimension reduction. A shared nearest neighbor network was created based on Euclidean distances between cells in multidimensional PC space (the first 50 PC were used) and a fixed number of neighbors per cell (50 neighbors). This was used to generate a two-dimensional Uniform Manifold Approximation and Projection (UMAP) for visualization as a harmonized atlas to dissect the cell populations from different treatment groups.

#### Differentially expressed genes

DEGs were calculated with “FindAllMarkers” function of Seurat in R 4.2 to study the different expression profiles in different cell types and up-regulated genes in different treatment groups (*47*). The “data” matrices of “SCT” assay were used, and the minimal log fold change was set to 0.25. Only genes that were detected in more than 25% of cells in either of the two populations were used to compute the DEGs with the Wilcoxon rank-sum test. Markers were identified as genes exhibiting significant up-regulation when compared against all other clusters and defined by having a Bonferroni-adjusted P-value < 0.05. The DEGs are ranked by the adjusted P-value to select the top DEGs for downstream analysis.

#### Cell type identification

To identify cell types, “FindClusters” function with the Leiden algorithm with the various resolution from 0.5 to 2.0 in the Seurat package were used for clustering (*47*). “FindAllMarkers” function was then applied. The top DEGs of individual clusters were examined for well-studied marker genes across literature and the clusters were then annotated for the most likely identity. For re-clustered IMs subsets, “FindClusters” was performed with different resolution until the right resolution is reached so that each cluster has unique gene expression pattern. Ten distinct cell types were identified, including IMs, non-classic IMs, recruited macrophages, non-classic monocytes, DC1, CD301b^-^.DC2, CD301b^+^.DC2, inflammatory DC2, migratory DCs, and cycling cells.

#### Gene ontology analysis

The gene set functional analysis was conducted with TopGO (*50*) (Fisher’s exact test) to get a general overview of the more represented functional categories with the DEGs in each cluster. Background genes of gene enrichment assays were defined as all the detected genes in datasets and the biological process was selected as the target of ontology analysis. The DEGs of different cell types were used as the enrichment input. Top 5 GO terms ranked by P-value (*51*) for every cell type was selected and compared with other cell types. Overlaps of those GO terms were removed from the final visualization.

#### Regulon analysis

The regulon specificity of chemokine-expressing IMs was analyzed using SCENIC package of R 4.2 and pySCENIC of Python 3.8 for the IM subtypes with count matrices as the input (*36, 37, 52*). The regulons and transcription factor activity (area under the curve) for each cell were calculated with motif collection version v10 as well as mm10_500bp_up_100bp_down_full_tx_v10_clust.genes_vs_motifs.rankings.feather and mm10_10kbp_up_10kbp_down_full_tx_v10_clust.genes_vs_motifs.rankings.feather databases from the cisTarget (https://resources.aertslab.org/cistarget/). The transcription factor regulons prediction was performed using the pySCENIC default parameters (*36, 37, 52*). The resulting AUC scores per each cell and adjacency matrices were used for downstream quantification of regulon specificity score. Top 10 transcription factor ranked by specificity score for every cell type was selected and compared with other cell types. Overlaps of those transcription factors were removed from the final visualization.

#### Pseudotime analysis

The package Monocle3 was performed to analyze single-cell trajectories (*53–56*). Pseudotime values were determined for all IMs. Monocle3 was used to learn the sequence of gene expression changes each cell must go through as part of a dynamic biological process and then arrange each cell in the trajectory (*53–56*). One node of each IM subset were set as the start allowing the visualization of the pseudotime process of naive IMs to activated IMs. To investigate the dynamic gene expression, DEGs over the pseudotime were calculated by the “graph_test” function. By ranking the P-value, top 100 pseudotime DEGs were identified and then visualized and the top 10 were labeled.

### scRNA-seq analysis of other tissues

The methods are available in GEO, accession code GSE193782, GSE215299, and SuperSeries GSE225668.

### Statistical Analysis

Statistical analysis was conducted using Prism software (GraphPad Software, Inc., San Diego, CA). All bar graphs are expressed as the mean (±SEM). Given the non-parametric nature of data requiring statistical analyses, we assessed statistical significance by performing two-tailed un-paired Mann-Whitney U tests. A P value less than 0.05 was considered statistically significant.

## Supplementary Materials

Fig. S1. Distinct transcriptional profiles and gene ontology enrichment observed in macrophage subsets.

Fig. S2. Absence of unique growth factor or M1/M2 gene signature expression in IM populations.

Fig. S3. M1/M2 phenotype analysis in IMs.

Fig. S4. Unbiased clustering analysis with higher resolution shows chemokine gene enrichment in IM subtypes.

Fig. S5. Chemokine-expressing IM subsets are conserved across multiple organs and species.

Fig. S6. Chemokine-expressing IM subsets are conserved in mouse heart (GSE179276).

Fig. S7. SCENIC analysis identifies regulons associated with chemokine regulation in different mouse lung IM subtypes, with the motif enriched 10k bp around the targeted gene TSS.

Fig. S8. SCENIC analysis identifies all regulons associated with chemokine regulation in different mouse lung IM subtypes, with the motif enriched 500 bp upstream to 100 bp downstream of the targeted gene TSS.

Fig. S9. SCENIC analysis identifies all regulons associated with chemokine regulation in different mouse lung IM subtypes, with the motif enriched 10k bp around the targeted gene TSS.

Fig. S10. Tissue-specific macrophages also exhibit chemokine-expressing subsets.

Fig. S11. Identification of the distinct subtypes of tissue-specific chemokine-expressing macrophages across different tissues and species.

Fig. S12. Chemokine-expressing monocyte subsets are conserved across multiple organs and species.

Fig. S13. IMs are required during iBALT formation. A. Representative IHC images for lung histopathological scoring for IM depleted mice in the HDM model compared to control. One set of three independent studies.

Fig. S14. Resembling cytokine-producing helper T cells, the specific combination of chemokines expresses by macrophages might also be tightly regulated, as proposed in the Results and Discussion sections.

## Acknowledgments

We are grateful to Dr. Wendy Wells, Steven H Boyle, Nichole R Lemelin, Jordan N Ray, Shannon N Schutz, Arnold J Wood, and Peter P Seery at the Department of Pathology at Dartmouth-Hitchcock Medical Center for kindly providing us with human breast reduction skin samples. RS-TS.

## Funding

National Institutes of Health grants R01 HL115334 (CVJ)

National Institutes of Health grants R01 HL135001 (CVJ)

National Institutes of Health grants R35 HL155458 (CVJ)

National Institutes of Health grants T32AI007363 (ABM)

UGent grant BOF.MET.2021.0007.01 (Contract Nr: 01M01521) (NG)

National Cancer Institute Cancer Center Support Grant 5P30CA023108

National Institutes of Health S10 1S10OD030242

National Institutes of Health NIGMS P20GM130454

National Institutes of Health NIH S10 S10OD025235

## Author contributions

Conceptualization: XL, CVJ

Methodology: XL, ABM, SM, KR, WTK, FWK, NG, CVJ

Investigation: XL, ABM, SM, WTK, FWK, KR

Visualization: XL, CVJ, ABM

Funding acquisition: CVJ, ABM, FWK

Writing – original draft: XL, CVJ

Writing – review & editing: XL, CVJ, ABM, NG

## Data and materials availability

The sequencing raw data and processed data used in this article are available in GEO, accession code GSE193782 (human BAL), GSE215299 (mouse lung-draining LN), and SuperSeries GSE225668 (mouse lung and all the others). The processed data are also available for online visualization at https://ams-supercluster.cells.ucsc.edu, https://cells.ucsc.edu/?ds=ln-mono-dc, and https://cells.ucsc.edu/?ds=lung-interstitial-macrophage

