## Supplementary Figures for "Coordinated Chemokine Expression Defines Macrophage Subsets Across Tissues"

**Supplementary Figures**
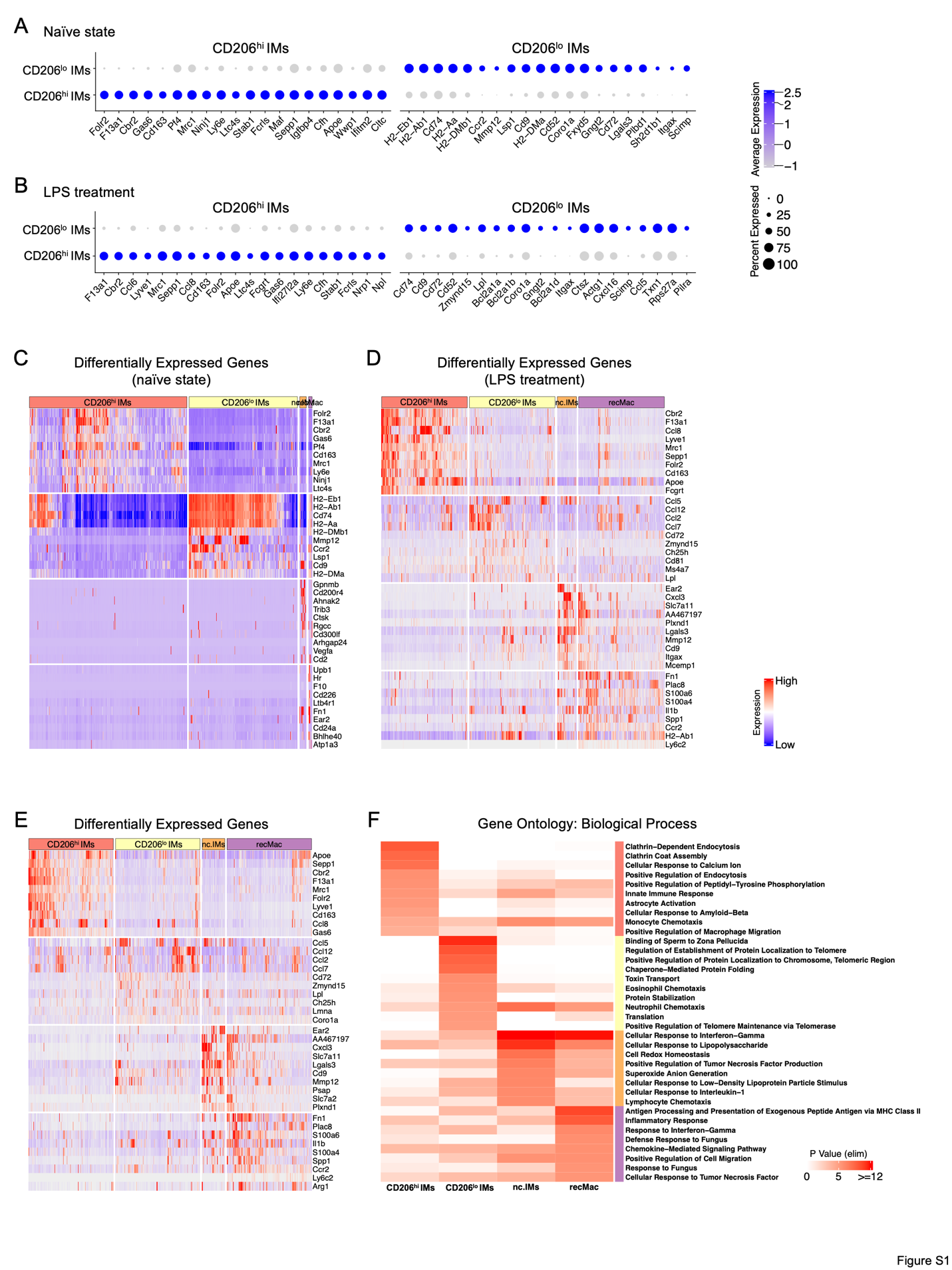

**Fig. S1. Distinct transcr­­­­­­­­­iptional profiles and gene ontology enrichment observed in macrophage subsets.** (**A**) Dot plot shows the expression of top 20 DEGs in CD206hi IMs and CD206lo IMs under naïve state. (**B**) Dot plot shows the expression of top 20 DEGs in CD206hi IMs and CD206lo IMs under LPS treatment. (**C**) Heat map shows the expression of top 10 DEGs in CD206hi IMs and CD206lo IMs, nc.IMs and rec.Mac under naïve state. (**D**) Heat map shows the expression of top 10 DEGs in CD206hi IMs and CD206lo IMs, nc.IMs and rec.Mac under LPS treatment. (**E**) Heat map shows the expression of top 10 overall DEGs in CD206hi IMs and CD206lo IMs, nc.IMs and rec.Mac. (F) Heat map shows the top 10 overall gene ontology terms (biological process) for CD206hi IMs and CD206lo IMs, nc.IMs and rec.Mac.

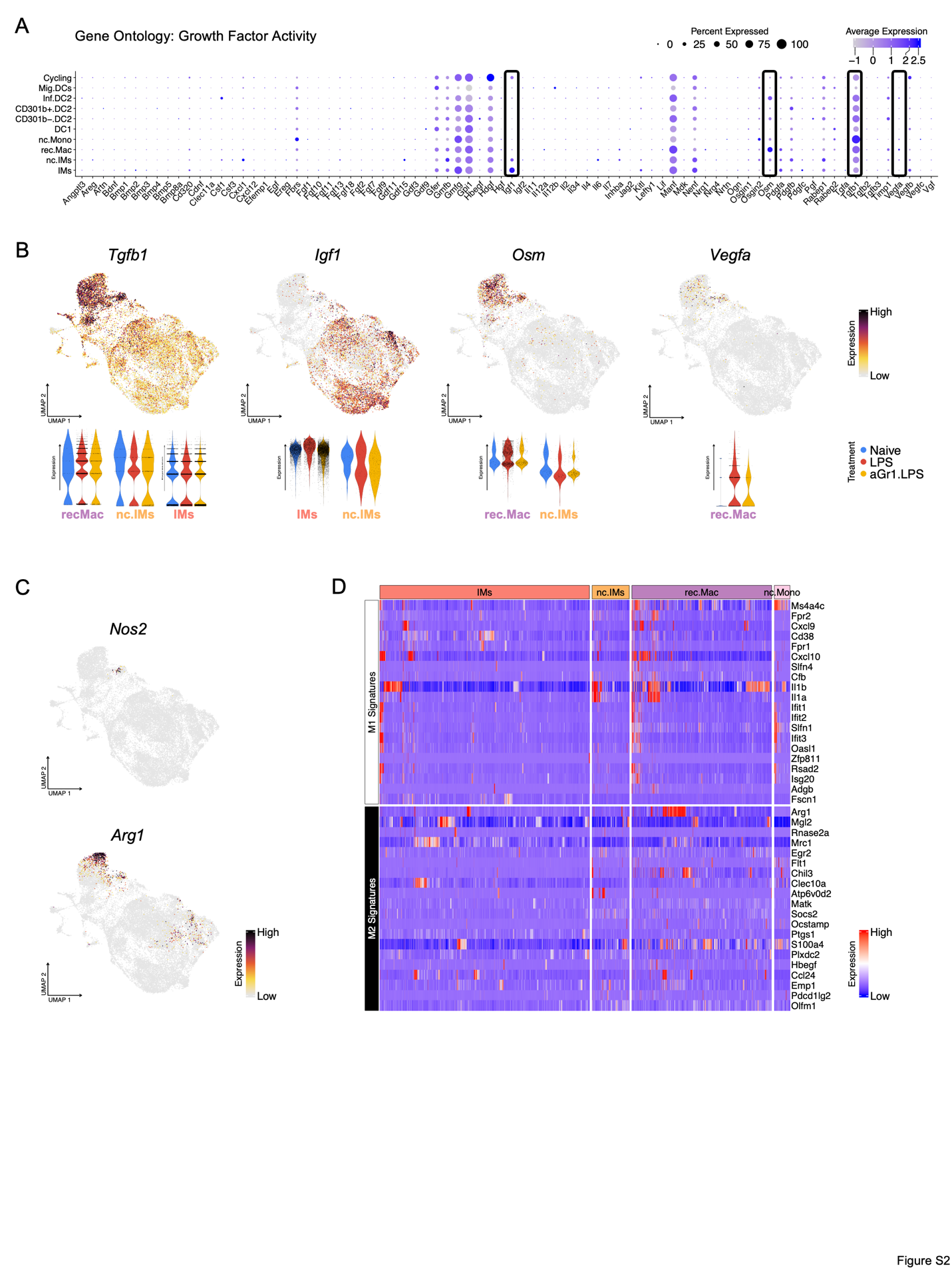

**Fig. S2. Absence of unique growth factor or M1/M2 gene signature expression in IM populations.** (**A**) Dot plot shows the expression of growth factor genes (GO: 0008083) in each individual cell type. (**B**) Feature plots show the expression of selected growth factor genes and violin plots show the expression of those genes in individual experimental groups. (**C**) Feature plots show the expression of Nos2 and Arg1, the markers for M1/M2 macrophages. (**D**) Heat map shows the expression of M1/M2 signature genes in CD206hi IMs and CD206lo IMs, nc.IMs and rec.Mac.

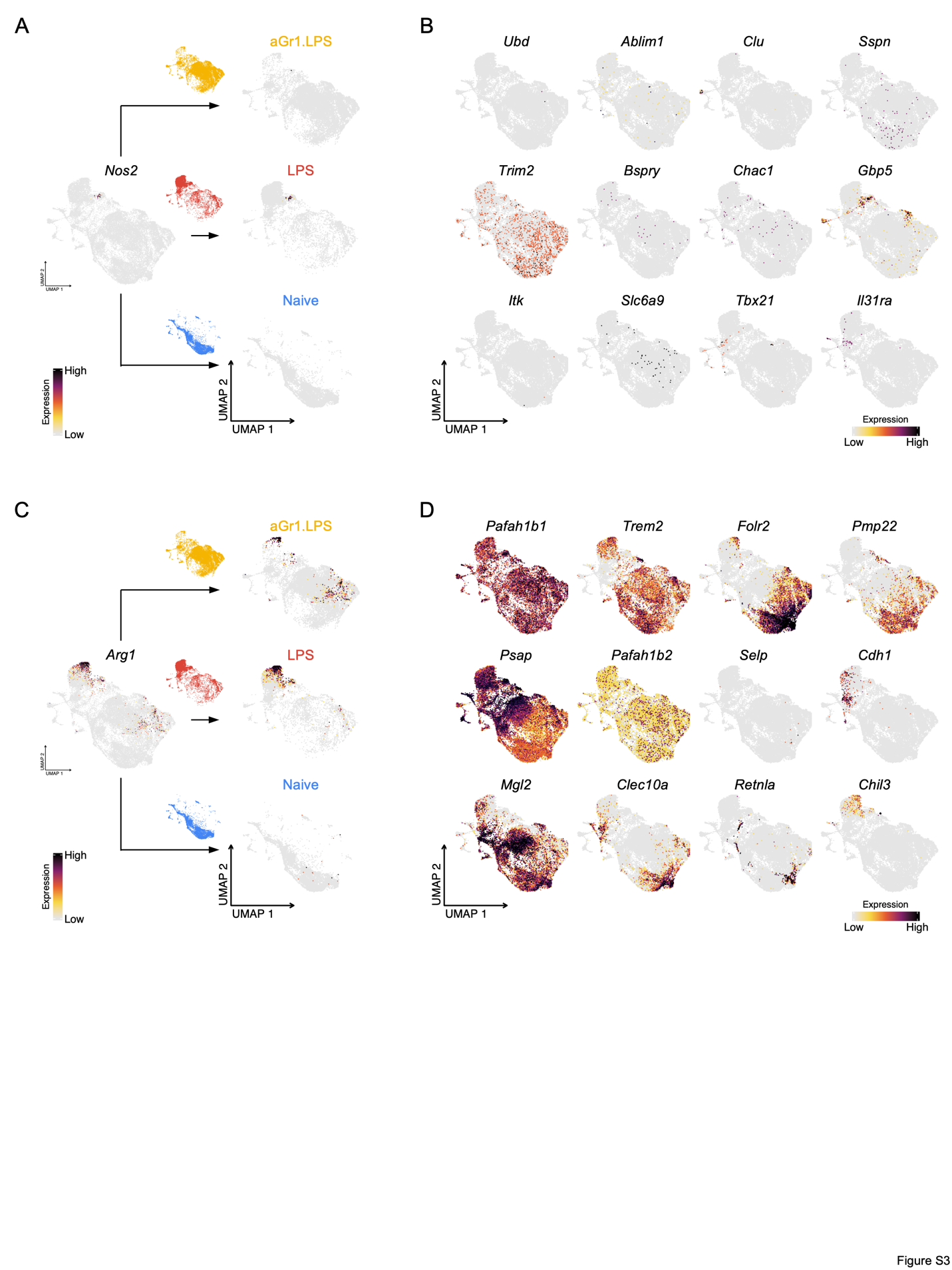

**Fig. S3. M1/M2 phenotype analysis in IMs.** (**A**) Feature plots show the expression of Nos2 in IMs in individual experimental groups. (**B**) Feature plots show the expression of M1 marker genes in IMs. (**C**) Feature plots show the expression of Arg1 in IMs in individual experimental groups. (**D**) Feature plots show the expression of M2 marker genes in IMs.

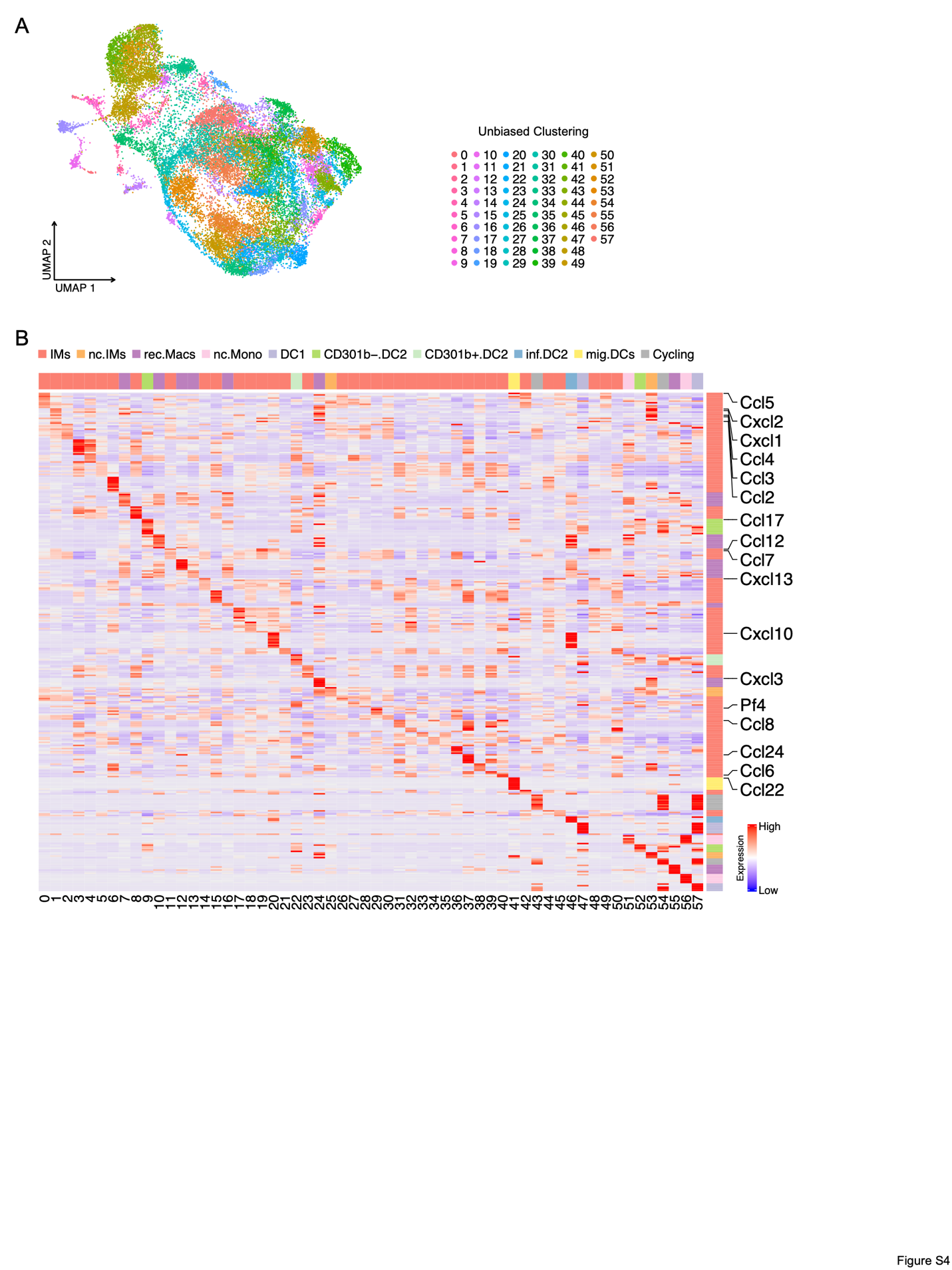

**Fig. S4. Unbiased clustering analysis with higher resolution shows chemokine gene enrichment in IM subtypes.** (**A**) UMAP plot shows the 58 clusters created by unbiased clustering analysis with higher resolution, including 38 IM clusters. (**B**) Heat map shows the expression of top 10 DEGs in each IM cluster. Chemokine genes are labeled.

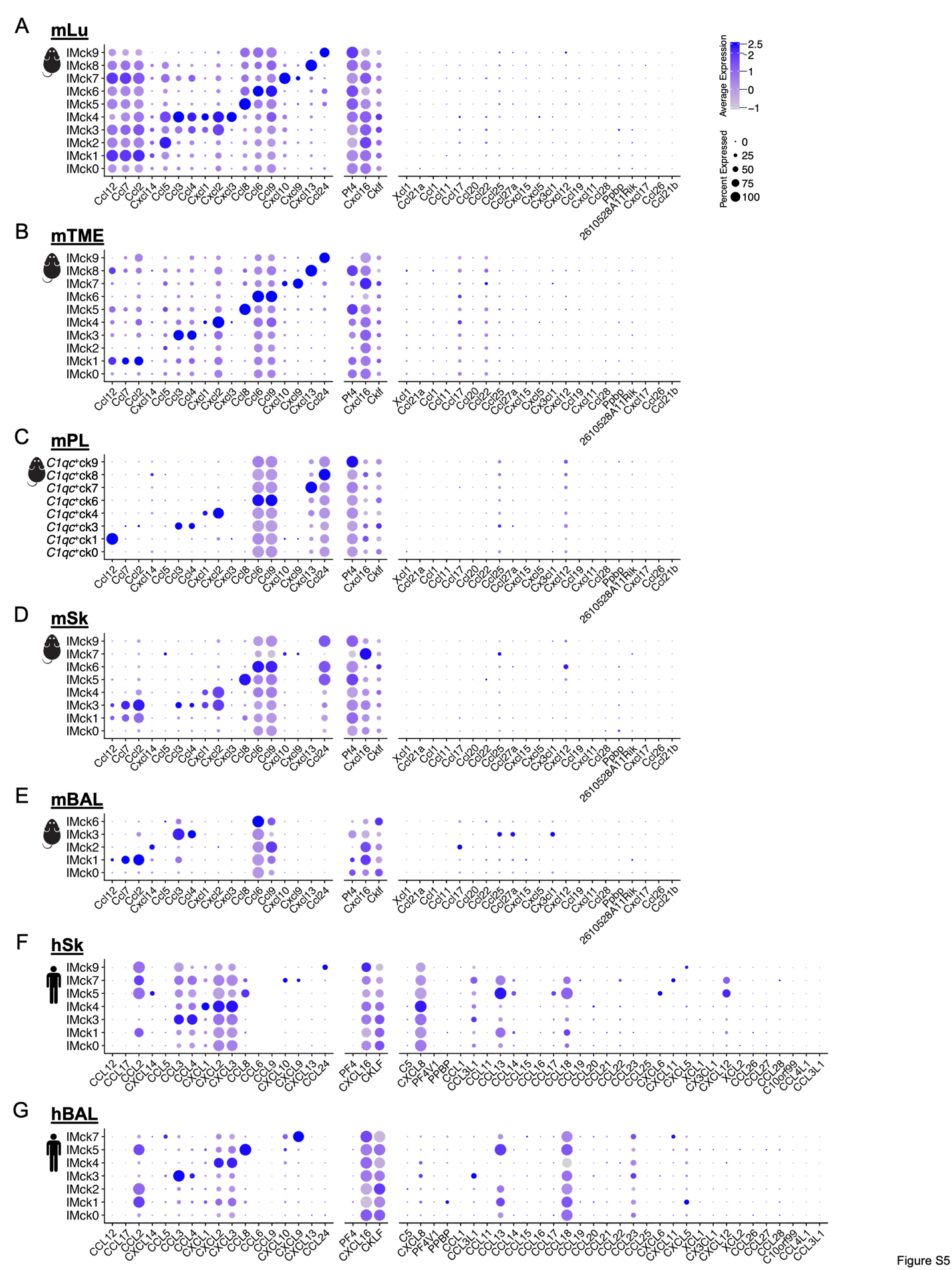

**Fig. S5. Chemokine-expressing IM subsets are conserved across multiple organs and species.** **A**-**G**. Dot plots show the expression of complete chemokine genes (GO: 0008009) in each IM chemokine-expressing subset in different tissues and species: (**A**) Mouse lung (mLu); (**B**) mTME; (**C**) mSk; (**D**) mBAL; (**E**) mPL; (**F**) hSk; (**G**) hBAL.

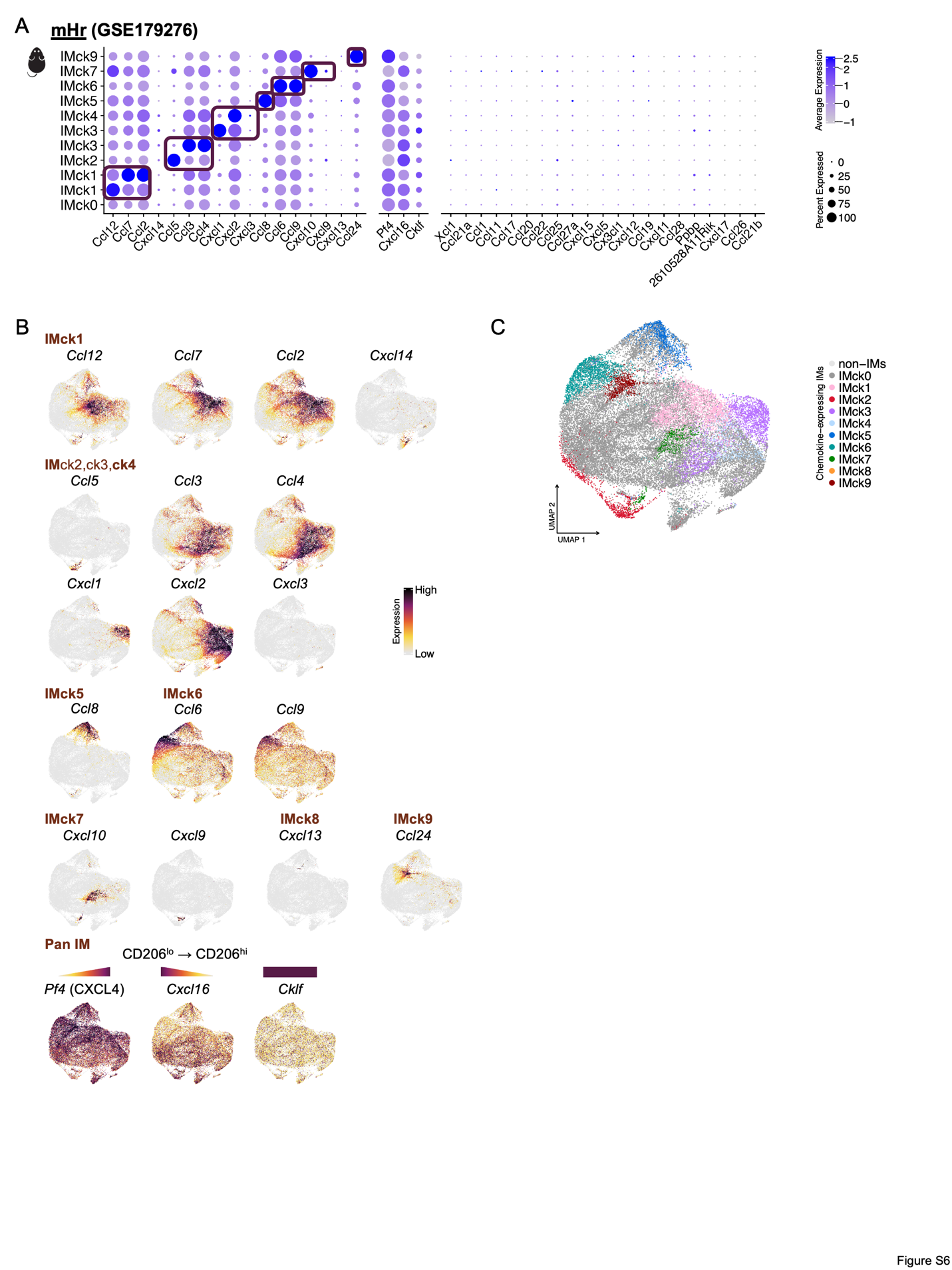

**Fig. S6. Chemokine-expressing IM subsets are conserved in mouse heart (GSE179276).** (**A**) Dot plot shows the expression of chemokine genes in each IM chemokine-expressing subset. (**B**) Feature plots show the expression of chemokine genes in each IM chemokine-expressing subset. (**C**) UMAP plot shows 10 IM subsets based on the expression levels of various chemokines (ck) : IMck0 to IMck9.

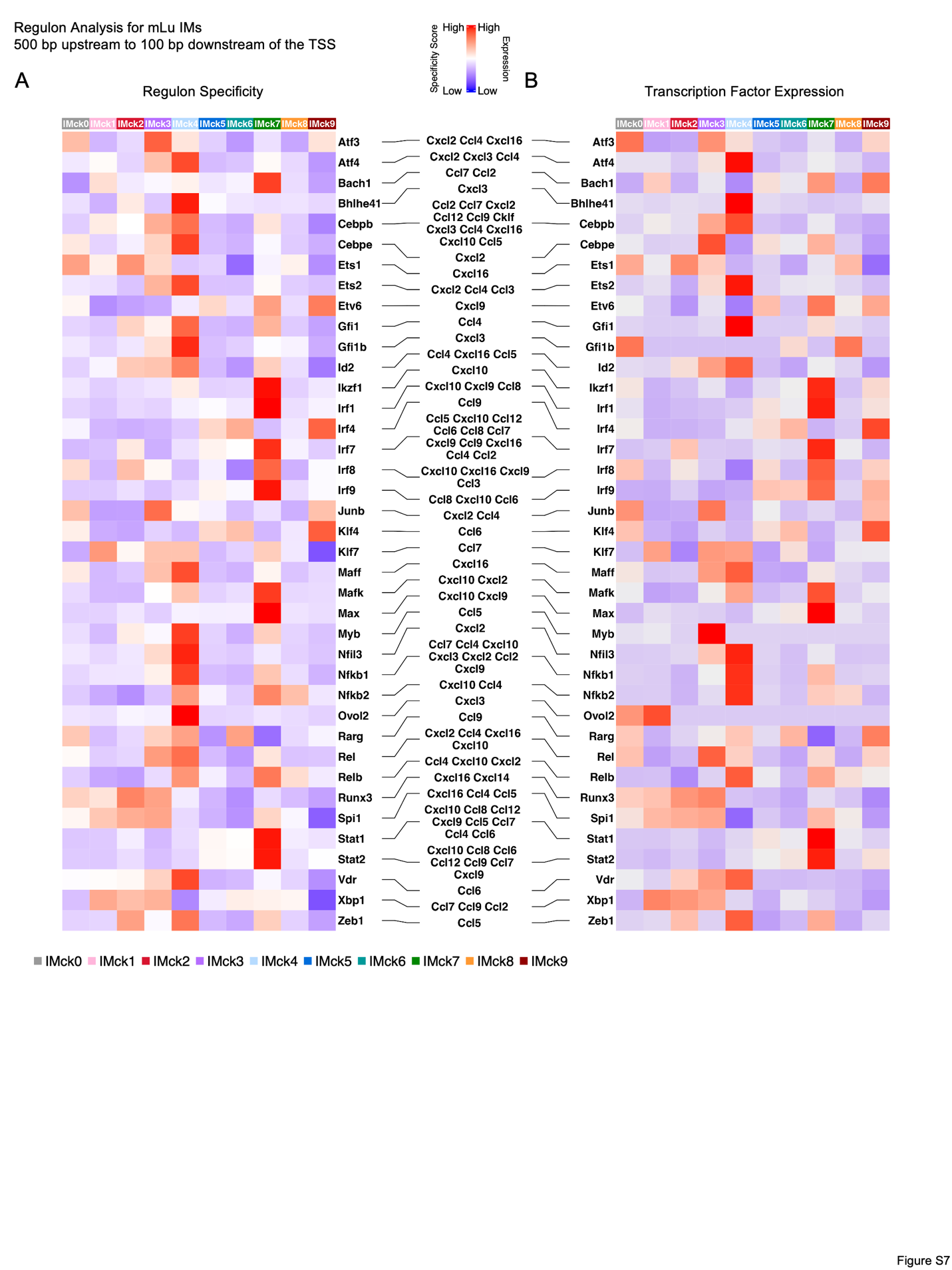

**Fig. S7. SCENIC analysis identifies regulons associated with chemokine regulation in different mouse lung IM subtypes, with the motif enriched 10k bp around the targeted gene TSS.** (**A**) Heat map shows the specificity of top 10 enriched regulons in each chemokine-secreting IM subtype, with the ones that putatively regulate the observed chemokine genes highlighted. (**B**) Heat map shows the expression of the transcription factors from the top 10 enriched regulons in each chemokine-secreting IM subtype, with the ones that putatively regulate the observed chemokine genes highlighted.

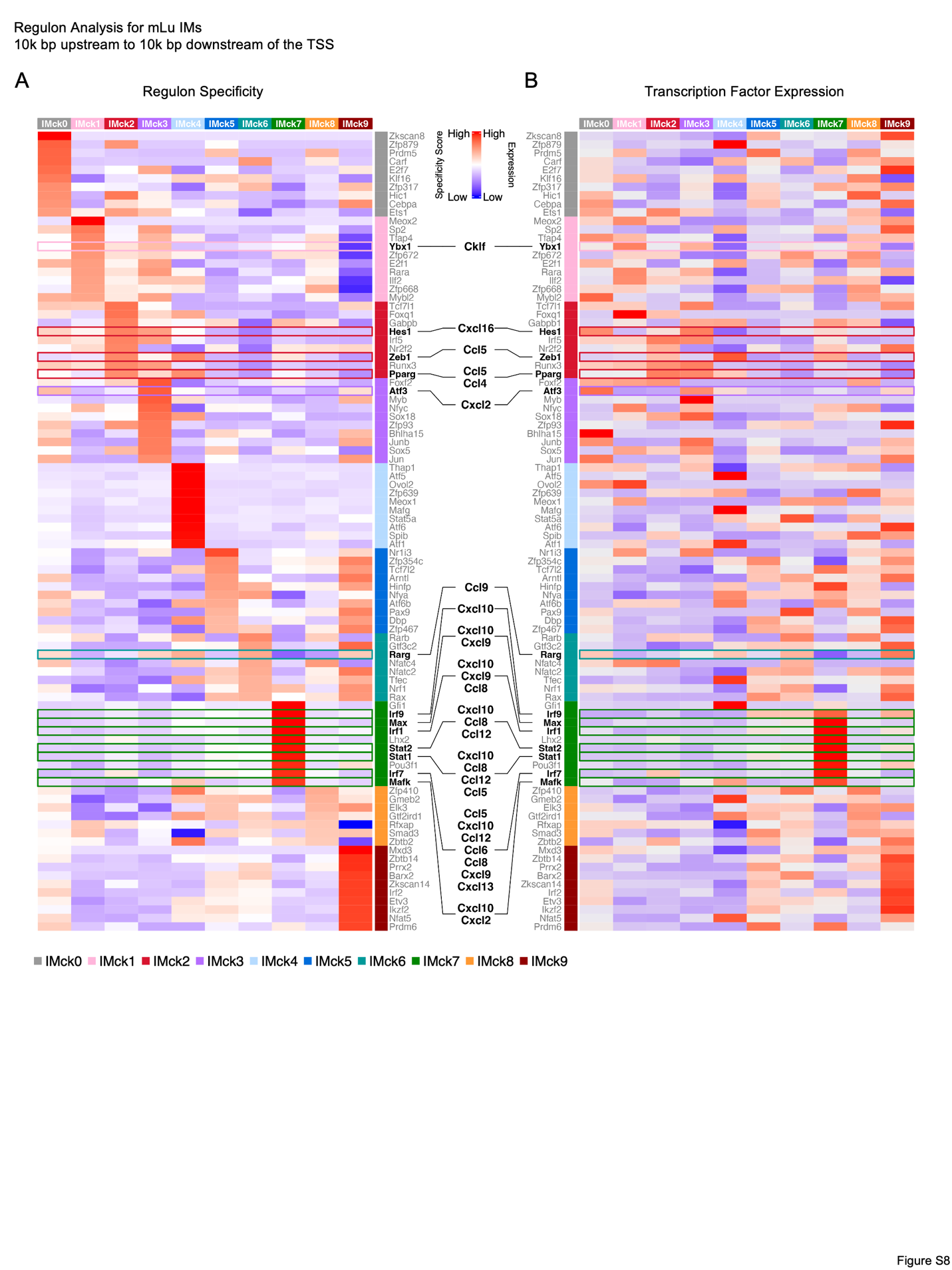

**Fig. S8. SCENIC analysis identifies all regulons associated with chemokine regulation in different mouse lung IM subtypes, with the motif enriched 500 bp upstream to 100 bp downstream of the targeted gene TSS.** (**A**) Heat map shows the specificity of all regulons associated with chemokine regulation in each chemokine-secreting IM subtype. (**B**) Heat map shows the expression of all transcription factors associated with chemokine regulation in each chemokine-secreting IM subtype.

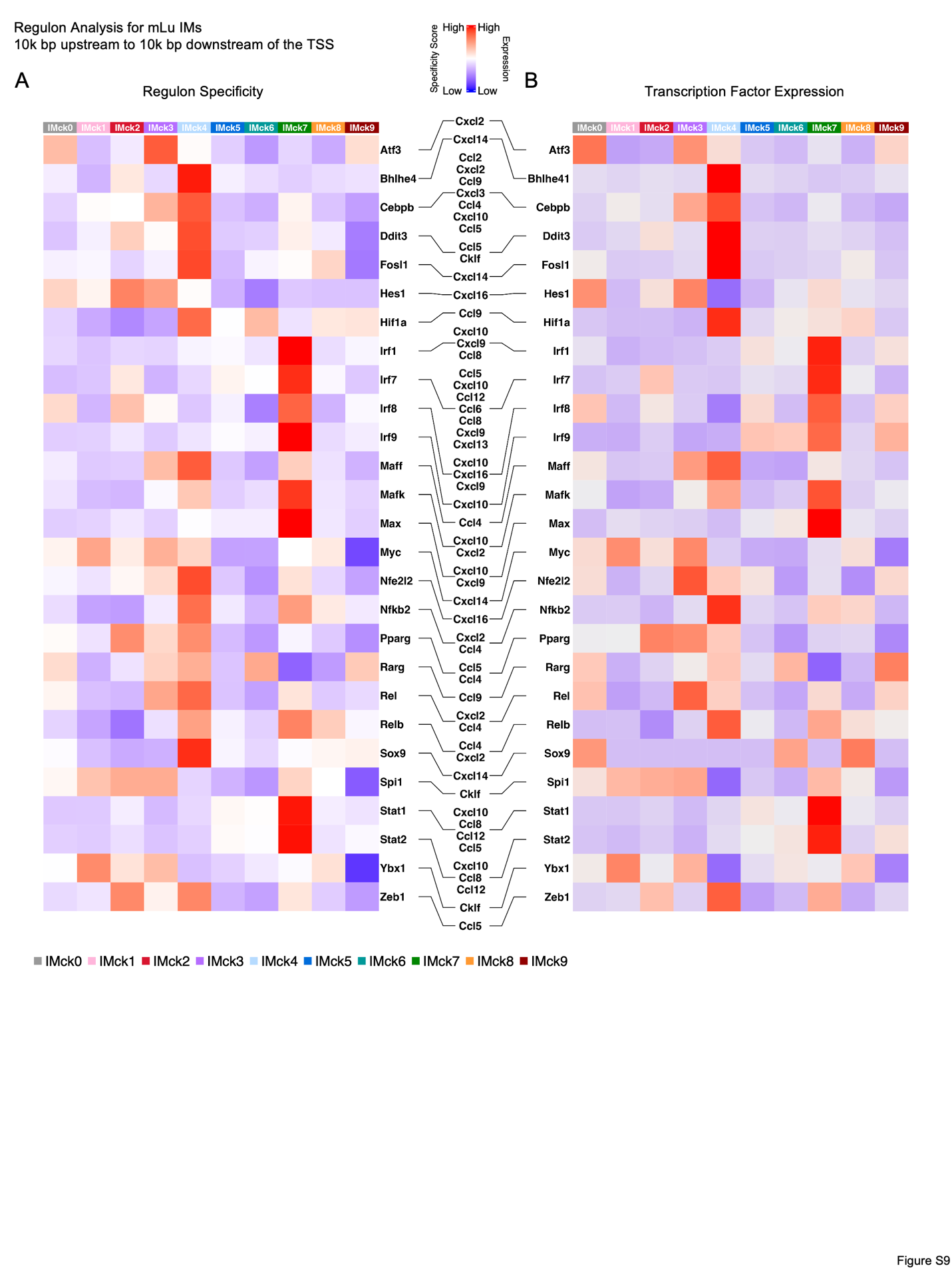

**Fig. S9. SCENIC analysis identifies all regulons associated with chemokine regulation in different mouse lung IM subtypes, with the motif enriched 10k bp around the targeted gene TSS.** (**A**) Heat map shows the specificity of all regulons associated with chemokine regulation in each chemokine-secreting IM subtype. (**B**) Heat map shows the expression of all transcription factors associated with chemokine regulation in each chemokine-secreting IM subtype.

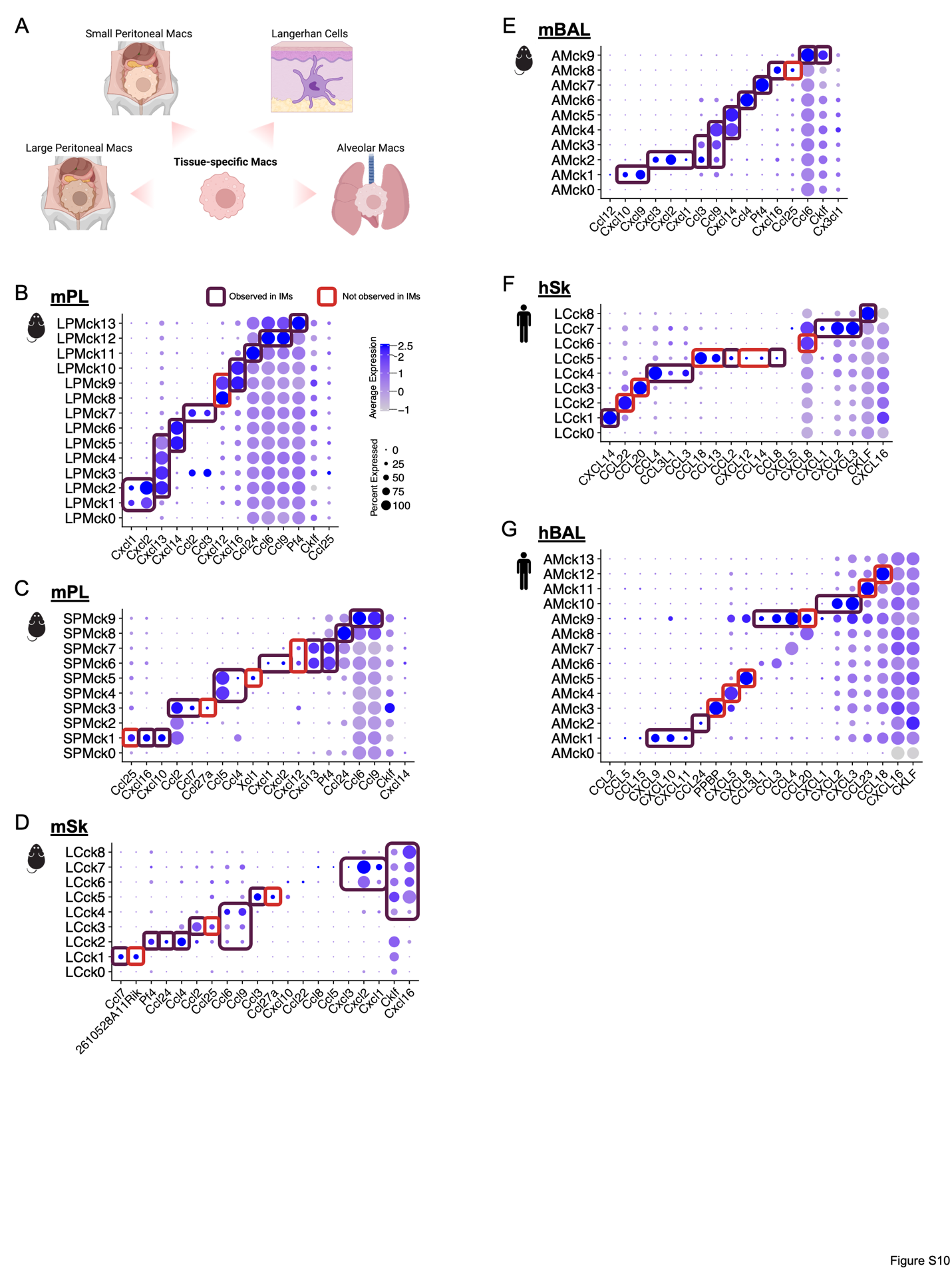

**Fig. S10. Tissue-specific macrophages also exhibit chemokine-expressing subsets.** (**A**) Diagram illustrates the origin of tissue-specific macrophages analyzed in this figure for chemokine expression. **B**-**G**. Dot plots show the expression of chemokine genes in each tissue-specific chemokine-expressing macrophage subsets in different tissues and species, with the chemokine combinations observed in IMs highlighted in dark boxes and the unique chemokines expressed by each macrophage population highlighted in red boxes. (**B**) Large peritoneal macrophages (LPMs) in mPL; (**C**) Small peritoneal macrophages (SPMs) in mPL; (**D**) Langerhans cells (LCs) in mSk; (**E**) Alveolar macrophages (AMs) in mBAL; (**F**) Langerhans cells (LCs) in hSk; (**G**) Alveolar macrophages (AMs) in hBAL.

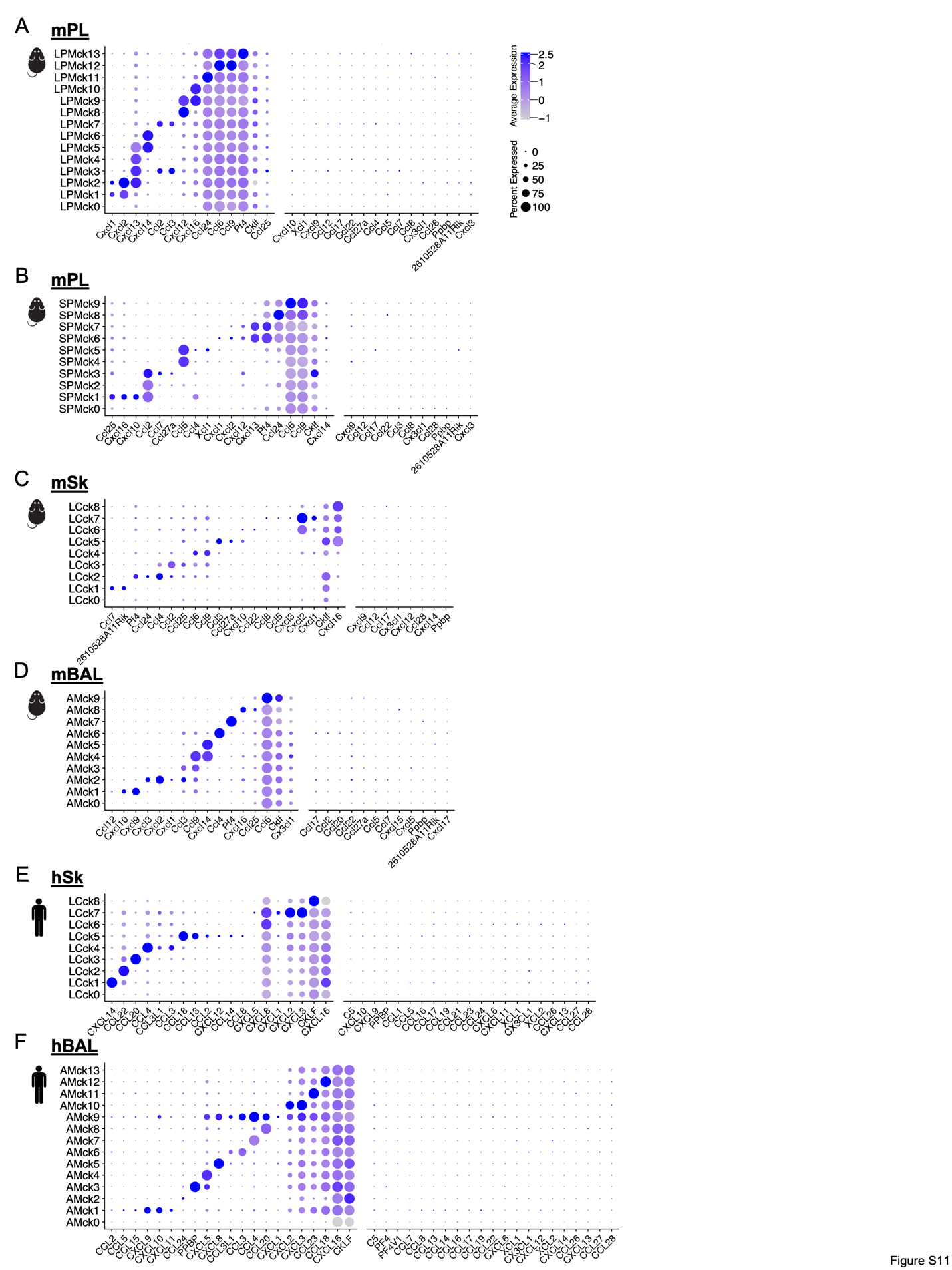

**Fig. S11. Identification of the distinct subtypes of tissue-specific chemokine-expressing macrophages across different tissues and species. A**-**F**. Dot plots show the expression of complete chemokine genes (GO: 0008009) in each tissue-specific chemokine-expressing macrophage subsets in different tissues and species. (**A**) LPMs in mPL; (**B**) SPMs in mPL; (**C**) LCs in mSk; (**D**) AMs in mBAL; (**E**) LCs in hSk; (**F**) AMs in hBAL.

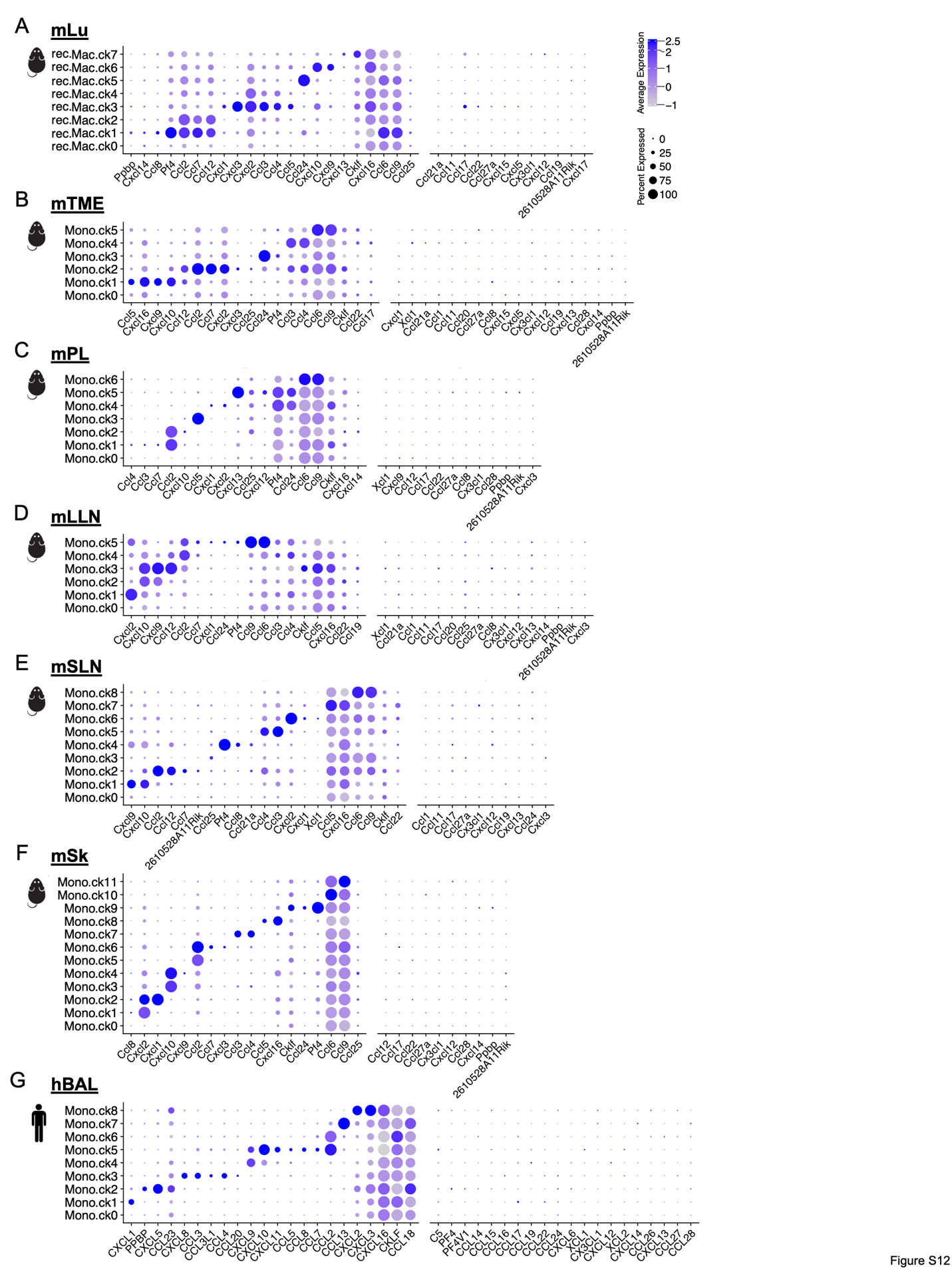

**Fig. S12. Chemokine-expressing monocyte subsets are conserved across multiple organs and species. A**-**G**. Dot plots show the expression of complete chemokine genes (GO: 0008009) in each chemokine-expressing monocyte subsets in different tissues and species. (**A**) mLu; (**B**) mTME; (**C**) mPL; (**D**) Mouse lung-draining lymph node (mLLN); (**E**) Mouse skin-draining lymph node (mSLN); (**F**) mSk; (**G**) hBAL.

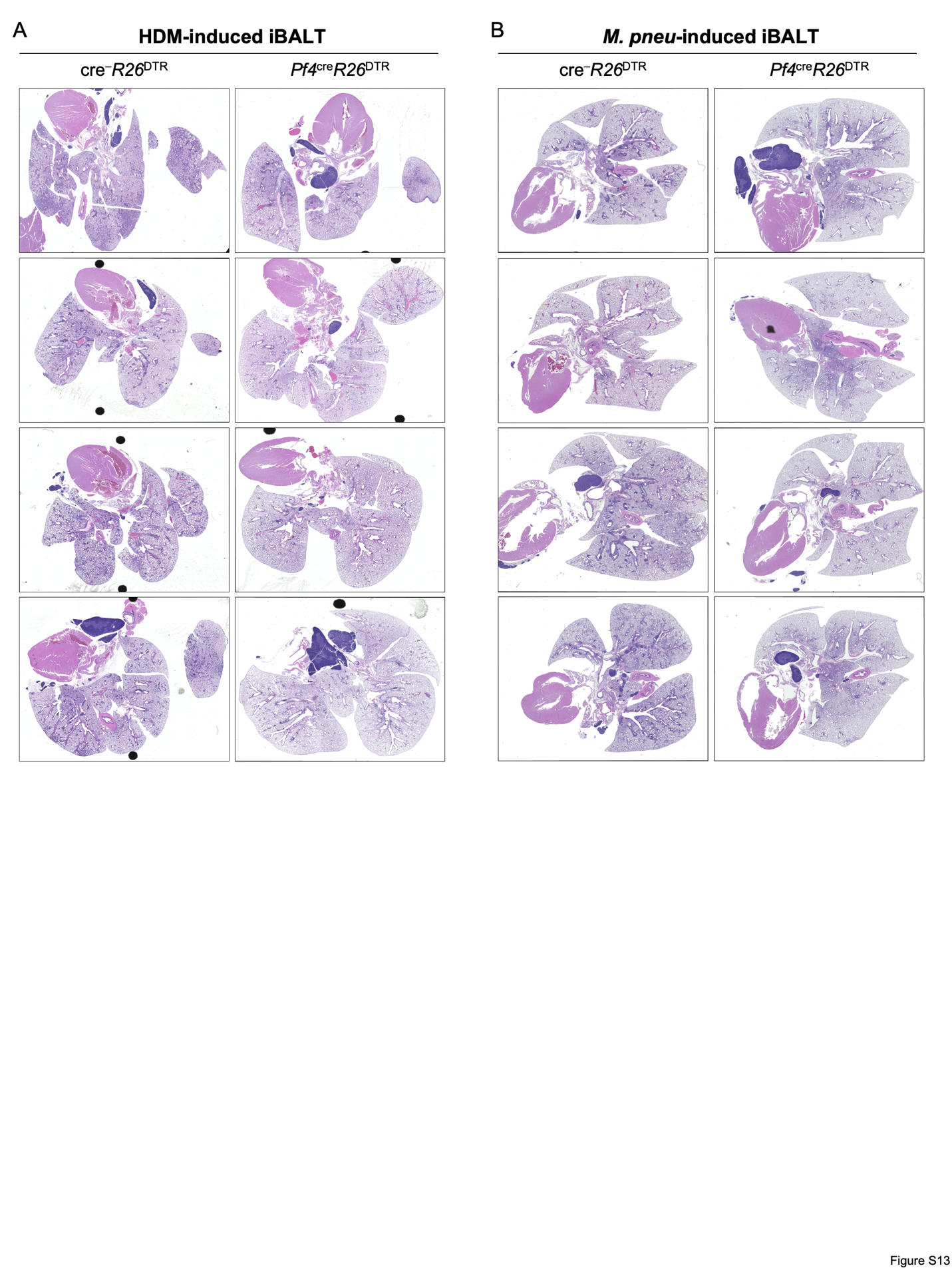

**Fig. S13. IMs are required during iBALT formation**. (**A**) Representative IHC images for lung histopathological scoring for IM depleted mice in the HDM model compared to control. One set of three independent studies. (**B**) Representative IHC images for lung histopathological scoring for IM depleted mice in the M. pneumoniae model compared to control. One set of three independent studies.

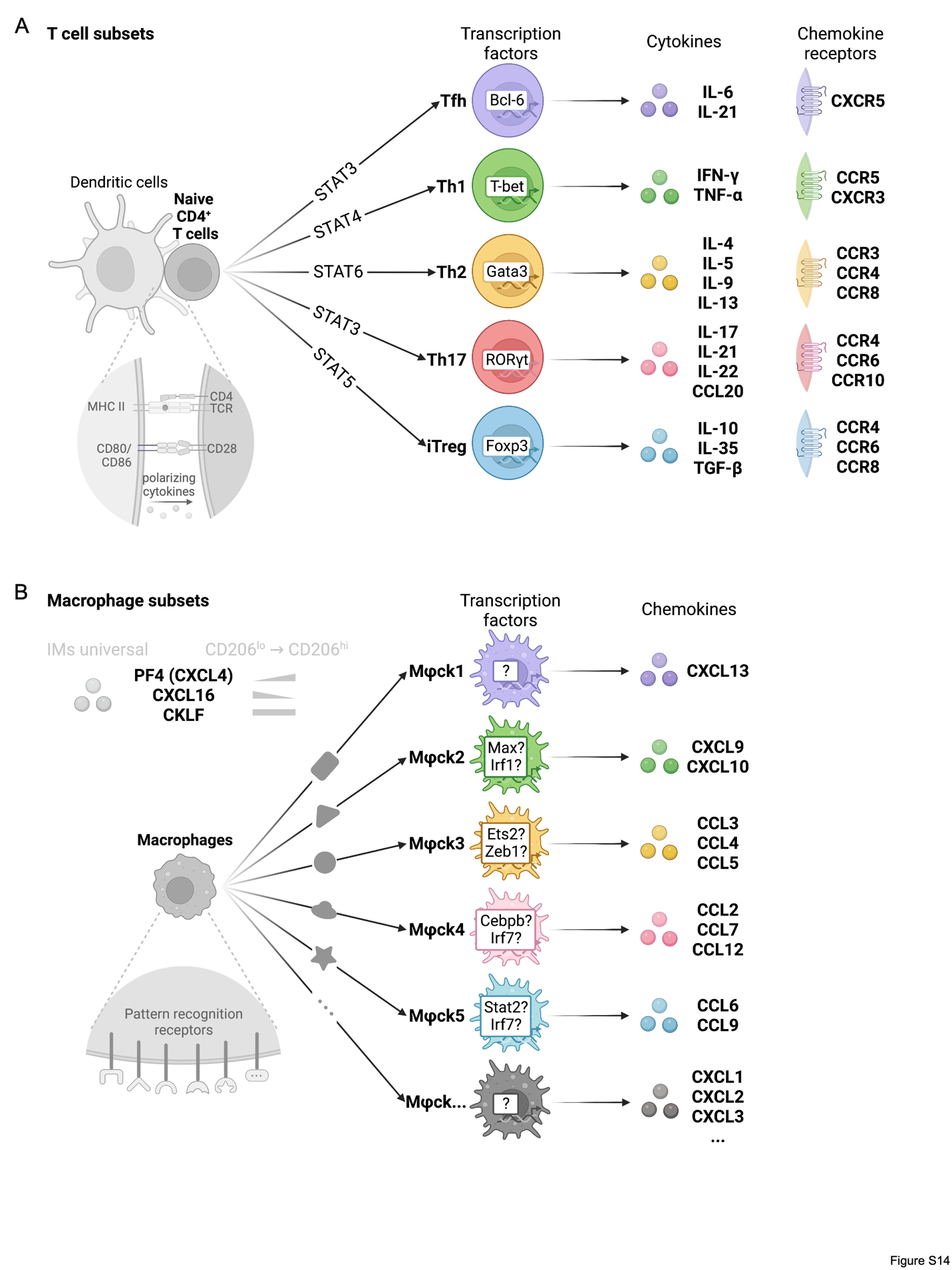

**Fig. S14. Resembling cytokine-producing helper T cells, the specific combination of chemokines expresses by macrophages might also be tightly regulated, as proposed in the Results and Discussion sections.** (**A**) The diagram summarizing the known universal T cell subtype biology, highlighting the specific differentiation routes, cytokines, and chemokine receptors expression patterns of different T cell subtypes. (**B**) The diagram summarizes the proposed universal macrophage subtype biology, highlighting the specific differentiation routes and chemokine expression patterns of different macrophage subtypes.
